# A stromal metabolic program suppresses NK-cell immunity to drive tumor progression in HER2-low breast cancer

**DOI:** 10.64898/2026.09.07.749862

**Authors:** Kayla Carter, Olajumoke Ogunlusi, Mrinmoy Sarkar, Stephen Akanbi, Christian Nguyen, Manasa Nekkanti, Akash Agarwal, Danielle Fails, Gayan I. Nawaratna, Cory Klemashevich, James Cai, Devon J. Boland, Tapasree Roy Sarkar

**Affiliations:** Department of Biology, Texas A&M University, College Station, TX, USA; Fortis Life Sciences, Montgomery, TX, USA; Department of Biochemistry and Biophysics, Texas A&M University, College Station, TX, USA; Integrated Metabolomics Analysis Core, Texas A&M University, TX, USA; Department of Veterinary Integrative Biosciences, Texas A&M University, TX, USA; Texas A&M Institute of Genome Sciences & Society (TIGSS), College Station, TX, USA; Texas A&M Center for Biological Clocks Research, College Station. TX, USA

## Abstract

Cancer-associated fibroblasts (CAFs) are major regulators of the tumor microenvironment, yet how distinct CAF states suppress innate immunity in HER2-low breast cancer remains poorly understood. Here, we identify an S100A4-enriched CAF population that expands during HER2-low breast tumor progression and establishes a metabolically immunosuppressive niche. Spatial transcriptomics and multiplex imaging of human HER2-low tumors reveal progressive CAF accumulation and an inverse spatial association between S100A4-enriched CAFs and immune infiltration, including natural killer (NK) cells. Using an immunocompetent HER2-low mammary tumor model, we show that S100A4-enriched CAFs promote tumor initiation and progression while suppressing NK-cell cytotoxicity, IFN-γ production, perforin, and granzyme B. Fractionation of CAF-conditioned media and metabolic profiling identify a low-molecular-weight immunosuppressive program characterized by enhanced branched-chain amino acid catabolism and accumulation of branched-chain α-keto acids (BCKAs). Mechanistically, BCKAs directly suppress NK-cell IFN-γ production, whereas inhibition of the branched-chain aminotransferase BCAT1 reduces CAF-mediated NK-cell suppression and restores antitumor cytotoxicity. BCAT1 inhibition also suppresses HER2-low tumor growth *in vivo*, an effect attenuated by NK-cell depletion, establishing NK-cell restoration as a functional component of its antitumor activity. Together, these findings uncover a CAF-driven metabolic immune checkpoint in which S100A4-enriched CAFs exploit BCAT1-dependent BCKA production to suppress NK-cell surveillance and promote HER2-low breast tumor progression. Targeting stromal BCAT1 therefore represents a potential strategy to dismantle CAF-mediated immune suppression and restore innate antitumor immunity.

## Introduction

Breast cancer is a biologically heterogeneous disease in which tumor behavior is shaped not only by malignant epithelial cells but also by dynamic interactions with the tumor microenvironment^1–3^. Among breast cancer subtypes, HER2-low tumors, defined by low but detectable ERBB2 expression without gene amplification, represent a large and clinically emerging category with distinct biology and therapeutic response^4–6^. While HER2-low breast cancers have gained attention due to the success of antibody–drug conjugates, the stromal and immune mechanisms that drive tumor progression in this subtype remain poorly defined^6,7^. Cancer-associated fibroblasts (CAFs) constitute one of the most abundant stromal populations in breast tumors and are now recognized as highly heterogeneous cells with context-dependent functions^8–11^. Rather than acting as passive structural elements, CAFs actively regulate tumor growth, immune evasion, and metabolic reprogramming through secreted factors, extracellular matrix remodeling, and metabolite exchange^12–15^. Single-cell and spatial transcriptomic studies have revealed that specific CAF subsets accumulate differentially across breast cancer subtypes and are linked to immunosuppressive microenvironments and poor clinical outcomes^9,16–18^. However, how defined CAF populations mechanistically shape anti-tumor immunity in HER2-low breast cancer remains largely unexplored.

Natural killer (NK) cells are critical mediators of early anti-tumor immune surveillance, exerting cytotoxic activity independently of antigen presentation^19,20^. Functional NK-cell infiltration is associated with improved clinical outcomes in breast cancer^21–23^, whereas Impaired NK-cell infiltration and function have been observed in breast tumors with poor immune responsiveness, yet the stromal mechanisms underlying NK-cell dysfunction are incompletely understood^24,25^. Emerging evidence suggests that CAFs can suppress NK-cell cytotoxicity. Accumulating evidence indicates that CAFs suppress immune cell function through both contact-dependent and paracrine mechanisms. Co-culture studies have shown that CAFs impair the cytotoxic activity of immune cells, including T cells and NK cells. In pancreatic ductal adenocarcinoma (PDAC), a Netrin G1–expressing CAF subset has been shown to inhibit NK-cell cytotoxicity and cytokine production^26^. Similarly, in melanoma, co-culture of NK cells with tumor-associated fibroblasts reduces NK-cell cytotoxicity, in part through prostaglandin E2 (PGE2)–mediated signaling^24^. In triple-negative breast cancer (TNBC), CAF–NK co-culture leads to potent downregulation of key activating receptors, including DNAX accessory molecule-1 (DNAM-1) and natural killer group 2D (NKG2D), thereby limiting NK-cell effector function^24^. Most studies have focused on CAF–NK co-culture, leaving paracrine mechanisms of NK-cell inhibition poorly understood, especially in breast cancer. Here, we identify a CAF-derived “paracrine suppressive circuit” mechanism that suppresses NK-cell cytotoxicity.

Additionally, we identify S100A4-enriched CAFs as a dominant stromal population in HER2-low breast tumors. S100A4-enriched CAFs have been linked to aggressive disease, immune suppression, and metabolic plasticity^27,28^, but their functional impact on innate immune cells has not been mechanistically resolved. We demonstrate that S100A4-enriched CAFs reprogram branched-chain amino acid (BCAA) metabolism by upregulating branched-chain aminotransferase 1 (BCAT1), leading to increased production of branched-chain keto acids (BCKAs) within the tumor microenvironment. Mechanistically, elevated BCKAs directly impair NK-cell cytolytic function, suppressing effector molecule expression and reducing tumor cell killing. This metabolite-driven immune suppression establishes a permissive niche for HER2-low tumor growth and progression. Notably, disruption of the S100A4–BCAT1–BCKA axis restores NK-cell cytotoxicity and limits tumorigenesis, revealing a previously unrecognized metabolic–immune checkpoint controlled by CAFs.

Analysis of human HER2-low breast tumors using multiplex immunostaining and spatial single-cell transcriptomics revealed that S100A4-enriched CAFs are progressively enriched during tumor progression and are associated with elevated BCAT1 expression in the stromal compartment. Strikingly, S100A4-high regions exhibited reduced NK-cell infiltration, indicating an inverse spatial relationship between CAF activation and innate immune presence. These data implicate S100A4-enriched CAFs as key regulators of the immune microenvironment and suggest a link between stromal metabolic reprogramming and suppression of NK-cell function in human tumors. Together, these findings uncover a CAF-driven metabolic mechanism of innate immune suppression that is selectively enriched in HER2-low breast cancer. Our study establishes S100A4-enriched CAFs as key regulators of NK-cell function through branched-chain amino acid metabolism and identifies BCAT1-dependent BCKA production as a targetable pathway linking stromal metabolism to immune evasion and tumor progression.

## Results

### 1. CAFs promote HER2-low mammary tumorigenesis

Several studies indicate that CAFs constitute one of the most abundant stromal cell populations in solid tumors and play critical roles in promoting malignant phenotypes. However, the functional and phenotypic significance of CAFs in HER2-low breast cancer, a clinically important yet understudied subtype, remains poorly defined. To address this gap, we systematically evaluated CAF heterogeneity in human HER2-low breast tumors across different stages of disease progression.

To define the spatial organization of the HER2-low tumor microenvironment during disease progression, we performed Xenium-based single-cell spatial transcriptomic analysis (**Fig 1A)** on human HER2-low breast tumor specimens (N=9) across stage I, stage II, and stage III disease (**Fig 1B**). Unsupervised cell-type annotation identified diverse epithelial, stromal, endothelial, myeloid, lymphoid, and CAF populations within the HER2-low tumor microenvironment. Comparison across tumor stages revealed progressive remodeling of the cellular ecosystem, with a marked increase in the CAF compartment in advanced-stage tumors **(Fig. 1B**). This increase in CAF abundance was accompanied by a relative reduction in immune-cell infiltration, suggesting that stromal expansion during HER2-low tumor progression is associated with an immune-excluded phenotype. Spatial mapping further demonstrated that advanced HER2-low tumors contained larger CAF-rich regions compared with early-stage tumors (**Fig. 1C**). Importantly, tumors or tumor regions with high CAF abundance showed reduced infiltration of immune cells, whereas CAF-low regions displayed greater immune-cell infiltration (**Fig. 1D).** These findings indicate that CAF accumulation is spatially associated with an immunosuppressive tumor microenvironment in HER2-low breast cancer.

**Figure 1:**
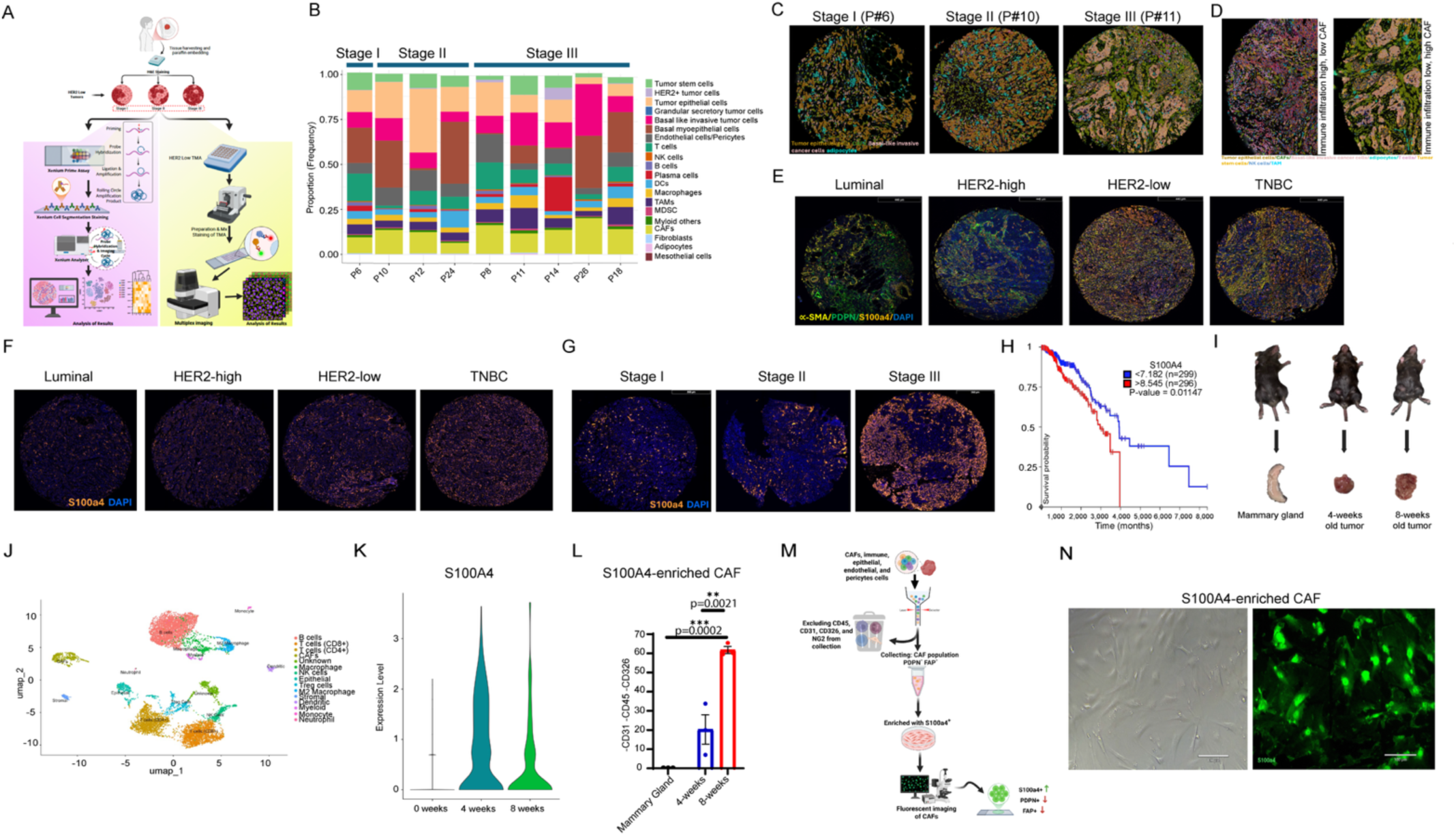
CAFs promote HER2-low tumorigenesis. **(A)** Schematic diagram showing the process of Xenium and multiplexed image acquisition. (**A**, Created with BioRender.com). **(B)** Comparison across tumor stages (n=9) reveals progressive remodeling of the cellular ecosystem. **(C)** Spatial mapping of HER2-low tumors containing larger CAF-rich regions compared with early-stage tumors. Scale bar: 1000µm. **(D)** CAF abundance in comparison to infiltration of immune cells. Scale bar: 1000µm. **(E)** Multiplex immunostaining of human breast tumor specimens (n=44) representing luminal, HER2-high, HER2-low, and triple-negative breast cancer (TNBC) subtypes. Scale bar: 440µm. **(F)** S100A4 was preferentially enriched in HER2-low and TNBC tumors in comparison to luminal and HER2-high subtypes. Scale bar: 350µm. **(G)** Immunostaining demonstrated that S100A4⁺ stromal cells increased from stage IA to stage IIIA. Scale bar: 350µm. **(H)** S100A4 was found to correlate with poor patient survival**. (I)** The mammary gland and tumors from 4- and 8-week-old C57BL/6J mice were harvested. **(J)** Analysis of 10x Genomics scRNA-seq exemplifies the tumor microenvironment, where **(K)** S100A4 is shown to be enhanced within the 4- and 8-week-old tumors. **(L)** Flow cytometric analysis showed a significant enrichment of the S100A4 population during tumor progression. Mean ± SEM; \*\**p* < 0.01, \*\*\**p* < 0.001 represent the significance level from a one-way ANOVA. (n=4-5 for each experimental condition) **(M)** Schematic diagram of Flow Activated Cell Sorting (FACS) exclusion of immune, epithelial, endothelial, and pericyte cells for the collection of NG2^-^PDPN^-^FAP^-^. (**M**, Created with BioRender.com). **(N)** FACS collection of S100A4-enriched CAFs. Scale bar: (Left) 120µm, (Right) 100µm. Indicated (n) represents the independent experiments as biological replicates.

Having observed progressive CAF expansion and immune exclusion in HER2-low tumors by single-cell spatial transcriptomics, we next asked whether this stromal remodeling reflected expansion of a specific CAF subtype. To address this, we performed multiplex immunostaining of human breast tumor specimens representing luminal, HER2-high, HER2-low, and triple-negative breast cancer (TNBC) subtypes and examined the expression of three major CAF-associated markers: α-SMA, PDPN, and S100A4^29^ (**Fig 1E).** This analysis revealed distinct subtype-associated CAF patterns. α-SMA was broadly detected across tumor subtypes without a marked subtype-specific enrichment, whereas PDPN expression was more prominent in luminal and HER2-high tumors. In contrast, S100A4 was preferentially enriched in HER2-low and TNBC tumors, suggesting that these more aggressive breast cancer subtypes are characterized by selective accumulation of S100A4-enriched CAFs rather than a uniform increase in all CAF populations (**Fig 1F**). We next examined S100A4 expression across progressive stages of HER2-low disease. Immunostaining demonstrated that S100A4-enriched stromal cells increased from stage IA to stage IIA and were most abundant in stage IIIA HER2-low tumors (**Fig. 1G**). This stage-associated increase in S100A4-enriched CAFs was consistent with the Xenium-based finding that CAF abundance increases during HER2-low tumor progression. Together, these human tumor data identify S100A4-enriched CAFs as a dominant stromal population enriched in HER2-low tumors and suggest that expansion of this CAF subset is associated with advanced disease and immune-excluded tumor architecture. By contrast, PDPN⁺ and FAP⁺CAF^30^ populations showed comparatively limited stage-dependent changes, indicating that HER2-low tumor progression is associated with selective expansion of the S100A4-enriched CAF compartment (**Supplementary Fig. 1A**). This pattern was consistent with the observation that S100A4-enriched CAFs increase during HER2-low tumor advancement and are most abundant in stage III tumors. Together, these human tumor data identify S100A4-enriched CAFs as a dominant stromal population enriched during HER2-low tumor progression. S100A4 was found to correlate with poor patient survival according to the TCGA database **(Fig. 1H**). Together, these data identify S100A4-enriched CAFs as a clinically relevant and dynamically expanding stromal population associated with tumor progression and adverse outcomes in human breast cancer.

To define CAF subtypes associated with HER2-low tumor progression, we next characterized stromal cell states in a murine model of HER2-low breast cancer. HER2 expression was readily detectable in MCF-7 and HER2-amplified SK-BR-3 cells, whereas Py230 and BT-20 cells exhibited comparatively low but measurable HER2 expression. In contrast, E0771, 4T1, and MDA-MB-231 cells showed minimal to undetectable HER2 expression. Based on this intermediate, low-level HER2 expression profile, Py230 was selected as a representative HER2-low model for our *in vivo* and mechanistic studies (**Supplementary Fig. 1B**). Py230 cells, previously established as a HER2-low model^31^, were orthotopically injected into the mammary fat pads of immunocompetent C57BL/6J mice. Tumors were harvested at key stages of development (4 and 8 weeks post-injection), and normal mammary fat pad fibroblasts (NMFs) from naïve mice (C57BL/6J) were collected as controls (**Fig. 1I**). Tumors and normal mammary fat pads were enzymatically dissociated into single-cell suspensions. Those cells were used for single-cell RNA-seq (scRNA-seq) using 10X Genomics. In total, we analyzed 8,987 high-quality single cells from four tumor-bearing mice and three naïve controls (**Fig. 1J**, **Supplementary Fig. 1C; Supplementary Table 1**). Using proper quality control (QC) and filtering, we obtained single-cell transcriptomic data for the studied tumors (N= 2/set). The tumor microenvironment was found to comprise diverse cell populations, including epithelial cells, endothelial cells, stromal cells, and multiple immune-cell populations, such as T cells, macrophages, natural killer cells (NK), and dendritic cells (DCs). ScRNA-seq analysis showed a significant enrichment of S100A4-enriched CAFs during tumor progression compared to PDPN⁺ and FAP⁺ CAF subsets (**Fig. 1K, Supplementary Fig. 1D, E**). This was further confirmed by flow cytometric analysis, which revealed a progressive increase in the S100A4-enriched CAF population during tumor progression (**Fig. 1L**). S100A4-enriched CAFs were subsequently isolated to assess their functional role in HER2-low tumorigenesis (**Fig. 1M**). To enrich for the S100A4⁺ CAF population, we first excluded major non-fibroblast lineages by sorting CD45⁻CD31⁻CD326/EpCAM⁻NG2⁻ cells, thereby removing immune cells, endothelial cells, epithelial/tumor epithelial cells, and pericyte-like cells^9^. Within this stromal fraction, we further excluded PDPN⁺ and FAP⁺ CAF subsets (**Fig.** 1N) and performed surface as well as intracellular staining^28^ for S100A4. The resulting CD45⁻CD31⁻CD326⁻NG2⁻PDPN⁻FAP⁻S100A4⁺ population was highly enriched for S100A4 expression and was operationally defined as the S100A4-enriched CAF population used in our functional and mechanistic studies (**Supplementary Fig. 2A, B**).

Normal mammary fibroblasts were isolated from enzymatically dissociated mouse mammary glands by FACS^32,33^. After exclusion of dead cells and lineage-positive immune, endothelial, epithelial, and pericyte-like cells using CD45, CD31, EpCAM/CD326, and NG2, respectively, fibroblasts were sorted from the live lineage-negative stromal fraction based on PDGFRα expression, as described previously^33^. The resulting live CD45⁻CD31⁻EpCAM/CD326⁻NG2⁻PDGFRα⁺ population was used as the normal mammary fibroblast fraction for downstream analyses (**Supplementary Fig. 2C**).

### 2. S100A4-enriched CAFs enhanced aggressive behavior of breast cancer cells

To comprehensively define how CAFs regulate tumor and immune cell behavior, we employed both S100A4-enriched CAF-conditioned media (CAF-CM) and direct S100A4-enriched CAF co-culture systems. These complementary approaches allow us to distinguish between paracrine (soluble factor– mediated)^34,35^ and contact-dependent or matrix-mediated mechanisms^36,37^. To examine the effects of CAFs derived from advanced-stage HER2-low tumors on breast cancer cells with distinct molecular subtypes, CAF-CM were collected from the S100A4-enriched CAFs isolated from 8-week-old tumors (**Fig. 2A**), as described in the Methods, and used to culture HER2-low (Py230, BT-20), triple-negative (MDA-MB-231, 4T1), and luminal (MCF-7) breast cancer cell lines. All tested cell lines exhibited significantly increased proliferation when cultured with CAF-CM compared with normal fibroblast–conditioned media (NF-CM) (**Fig. 2B**, **Supplementary Fig. 3A**). Functionally, CAF-CM markedly enhanced migratory capacity (**Fig. 2C, Supplementary Fig. 3B**) and mammosphere formation (**Fig. 2D, Supplementary Fig. 3C**) across cancer cell lines relative to NF-CM. Collectively, these findings indicate that CAF-secreted factors promote phenotypic plasticity and acquisition of aggressive, pro-metastatic traits in diverse breast cancer subtypes. Similarly, co-culture with S100A4-enriched CAFs recapitulated the phenotypes observed in tumor cells, indicating that CAF-mediated effects are driven by both paracrine and contact-dependent mechanisms (**Fig. 2E, F, Supplementary Fig. 4A, B**). To define the specific contribution of S100A4-enriched CAFs to these CAF-mediated effects, distinct CAF populations (FAP+, S100A4+, and PDPN+) were stained (**Supplementary Fig. 5A**) and additionally selectively inhibited in the CD45⁻CD31⁻CD326⁻NG2⁻ CAF population, individually, and conditioned media from the treated (and untreated) CAFs) were subsequently applied to breast cancer cells to assess changes in cellular behaviors. Selective inhibition of the FAP-enriched CAF population using UAMC1110 did not alter the pro-aggressive properties of CAF-conditioned media (CAF-CM), as cancer cells cultured with this CM continued to exhibit aggressive phenotypes comparable to those observed with untreated CAF-CM and significantly greater than cells cultured with normal fibroblast–conditioned media (NF-CM) (**Fig. 2G, Supplementary Fig. 5B**). Similarly, inhibition of the PDPN-enriched CAF population using gp38 InVivoMAb failed to attenuate CAF-CM–induced aggressive behaviors in cancer cells (**Fig. 2G, Supplementary Fig. 5B**). To define the contribution of S100A4-enriched CAFs to CAF-mediated tumor-promoting effects, we first examined the consequences of pharmacologically suppressing S100A4 using niclosamide, which has previously been shown to reduce S100A4 expression and signaling^38^. Treatment of CAFs with niclosamide reduced S100A4 expression and markedly attenuated the ability of CAF-CM to promote aggressive cellular phenotypes in HER2-low breast cancer cells, including mammosphere formation (**Fig. 2G; Supplementary Fig. 5B**). To independently validate the functional contribution of S100A4 to the CAF-mediated phenotype, we next used a genetic loss-of-function approach. Mouse-specific siRNA-mediated depletion of S100A4 in CAFs significantly reduced S100A4 expression and diminished the ability of CAF-conditioned media to enhance HER2-low breast cancer cell proliferation and mammosphere formation (**Fig. 2H**). Together with the pharmacologic studies, these findings establish S100A4 as an important component of the tumor-promoting program of S100A4-enriched CAFs.

**Figure 2.**
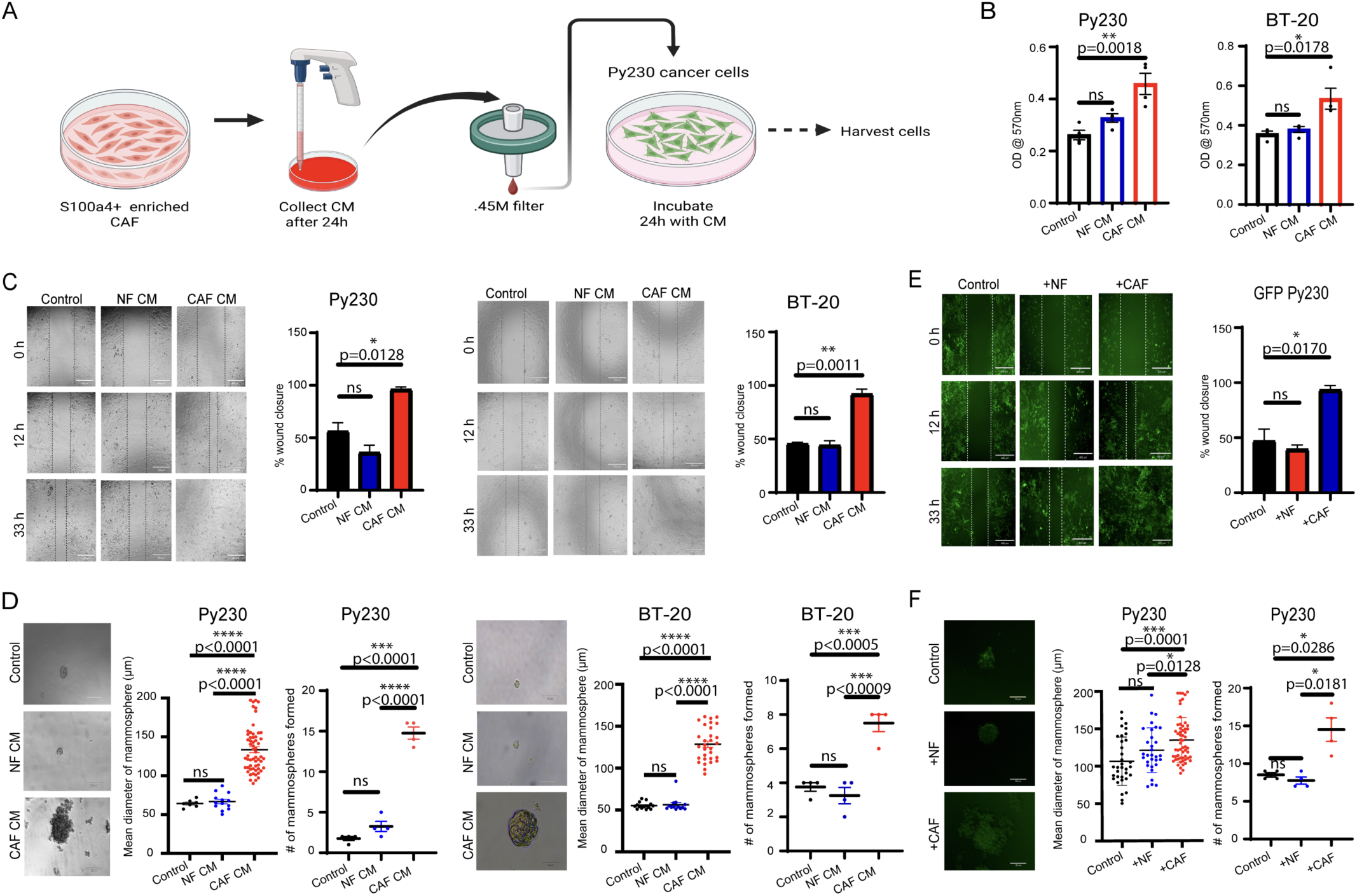
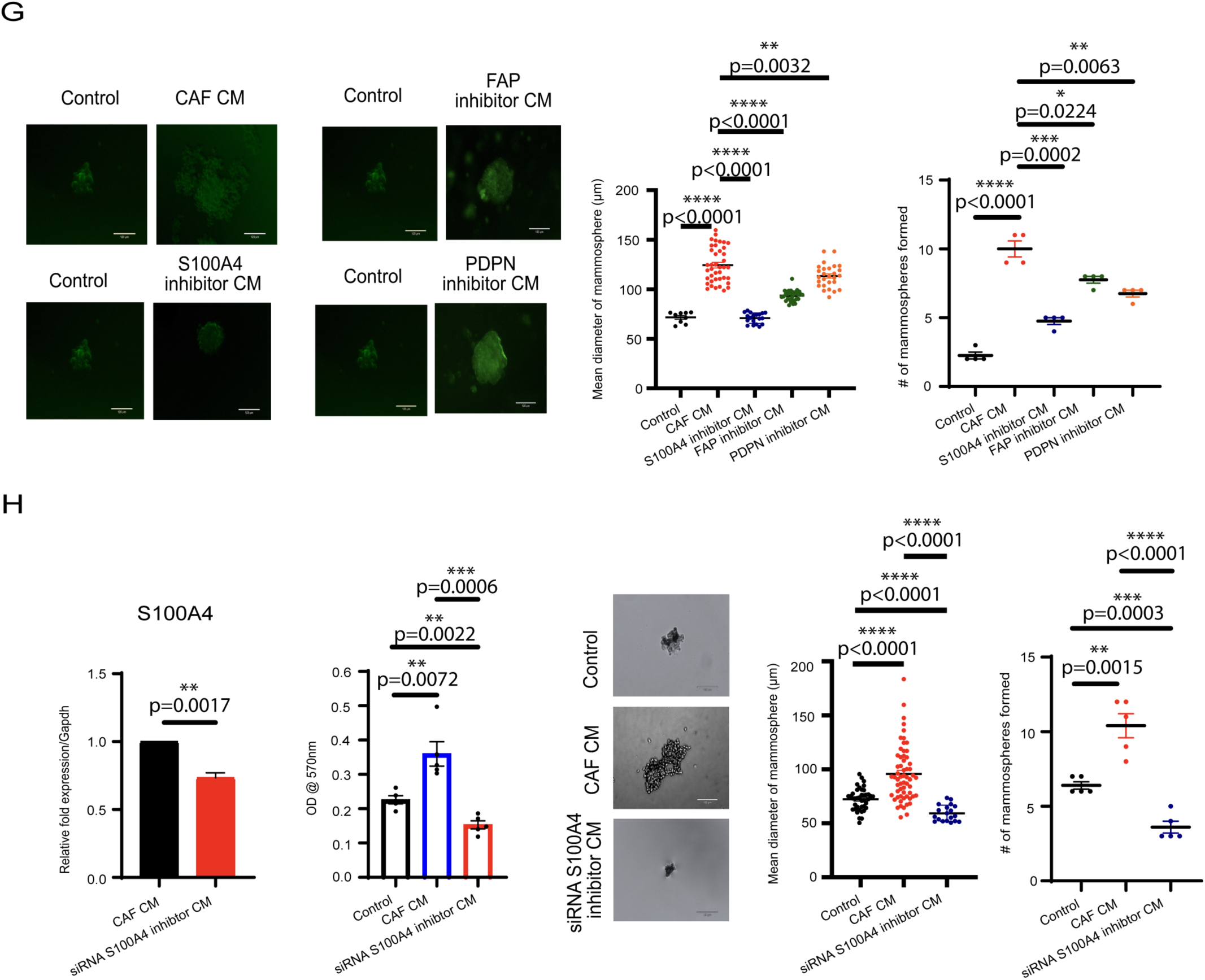
CAF CM and cells enhance the aggressive properties of HER2-low cancer cells. **(A)** Schematic diagram of conditioned media collection. (**A**, Created with BioRender.com). **(B)** MTT assay analyzing the cell proliferation of Py230 and BT-20 (n=4). Mean ± SEM; \**p* < 0.05, \*\**p* < 0.01 represent the significance level from a one-way ANOVA. **(C)** The addition of S100A4-enriched CAF CM compared to NF CM added or cells only for wound percent closure of Py230 and BT-20 cancer cells (n=4). Mean ± SEM; \**p* < 0.05, \*\**p* < 0.01 represent the significance level from a one-way ANOVA. Scale bar: 400µm. **(D)** Images along with mean diameter and number of mammospheres of Py230 and BT-20 (n=4). Mean ± SEM; \*\*\**p* < 0.001, \*\*\*\**p* < 0.0001 represent the significance level from a one-way ANOVA. Scale bar: 120µm. **(E)** The addition of S100A4-enriched CAF cells compared to NF cells or cells only for wound percent closure of GFP-tagged Py230 cancer cells (n=4). Mean ± SEM; \**p* < 0.05 represents the significance level from a one-way ANOVA. Scale bar: 400µm. **(F)** Images along with mean diameter and number of mammospheres of GFP-tagged Py230 cancer cells (n=4). Mean ± SEM; \**p* < 0.05, \*\*\**p* < 0.001 represent the significance level from a one-way ANOVA. Scale bar: 120µm. **(G)** Images along with the number of mammospheres of GFP-tagged Py230 cultured with S100A4-enriched CAF CM, S100A4-inhibited CM, PDPN-inhibited CM, and FAP-inhibited CM (n=4). Mean ± SEM; \**p* < 0.05, \*\**p* < 0.01, \*\*\**p* < 0.001, \*\*\*\**p* < 0.0001 represent the significance level from a one-way ANOVA. Scale bar: 120µm. **(H)** A mouse-specific siRNA was used to suppress S100A4 in CAFs, as shown by real-time PCR (n=4), MTT (n=5), and mammosphere experiments (n=5). PCR: Mean ± SEM; \**p* < 0.05, \*\**p* < 0.01 represent the significance level from an unpaired *t*-test. Assays: Mean ± SEM; \*\**p* < 0.01, \*\*\**p* < 0.001, \*\*\*\**p* < 0.0001 represent the significance level from a one-way ANOVA. Scale bar: 120µm. Indicated (n) represents the independent experiments as biological replicates.

Consistent with these *in vitro* findings, co-injection of Py230 cells with S100A4-enriched CAFs isolated from advanced-stage (8 weeks post-injection) HER2-low Py230 tumors resulted in a significant increase in tumor volume compared with Py230 cells co-injected with normal fibroblasts (NFs) *in vivo* (**Fig. 3A**). To determine whether pharmacologic suppression of S100A4-associated stromal activity affects HER2-low tumorigenesis *in vivo*, Py230 tumor-bearing mice were treated with niclosamide at distinct stages of tumor development (**Fig. 3B**) at tumor initiation, and 3 weeks post-implantation (to study the role in tumor progression). Niclosamide treatment initiated immediately following orthotopic Py230 cell transplantation into C57BL/6J mice markedly inhibited tumor initiation (**Fig. 3C, E**). Similarly, administration of niclosamide to established tumors at 3 weeks of age significantly suppressed tumor progression (**Fig. 3D, E**). These findings indicate that S100A4-enriched CAFs play a critical role in tumor initiation as well as in tumor progression. Niclosamide treatment inhibited epithelial-to-mesenchymal transition (EMT) in HER2-low tumors, as evidenced by reduced vimentin expression and increased E-cadherin expression in treated tumors (**Fig. 3F**).

**Figure 3.**
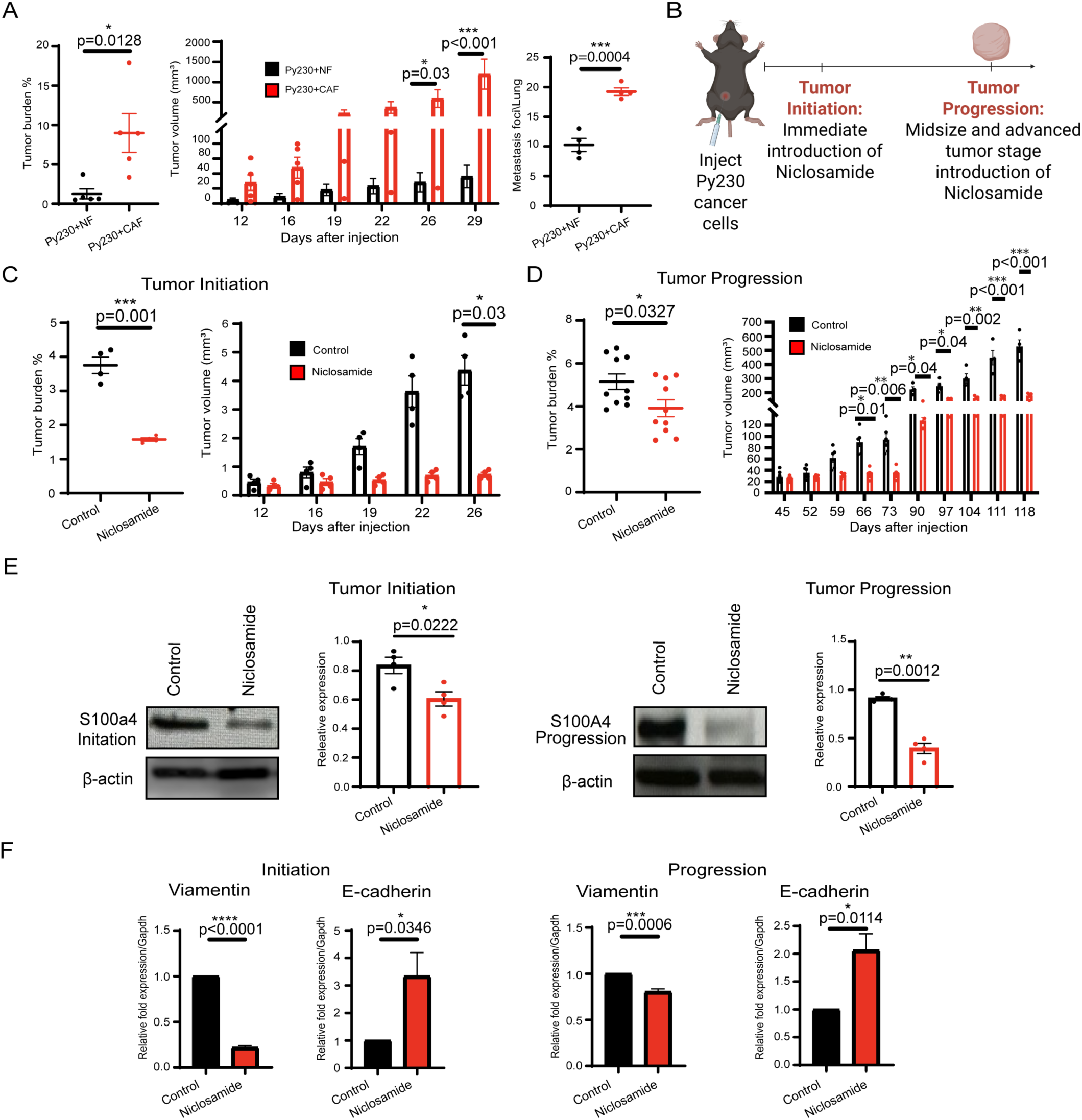
Pharmacologic suppression of S100A4-associated stromal activity inhibits HER2-low tumor initiation and progression. **(A)** HER2-low tumors harvested from C57BL/6J mice with co-injection of NF cells compared to S100A4-enriched CAF cells in tumor burden percentage (tumor-to-body-weight ratio), tumor volume, and lung metastasis (n=5 for each experimental condition). Tumor burden & lung metastasis: Mean ± SEM; \**p* < 0.05, \*\*\**p* < 0.001 represent the significance level from an unpaired two-sided *t*-test. Tumor volume: Mean ± SEM; \**p* < 0.05, \*\*\**p* < 0.001 represent the significance level from a one-way ANOVA. **(B)** Schematic diagram of niclosamide treatment via i.p. injection at different stages of tumor development. (Created with BioRender.com). Tumor burden percentage (tumor-to-body-weight ratio) and tumor volume (n=5 for each experimental condition), tumor initiation **(C),** and tumor progression **(D)**. Tumor burden: Mean ± SEM; \**p* < 0.05, \*\*\**p* < 0.001 represent the significance level from an unpaired two-sided *t*-test. Tumor volume: Mean ± SEM; \**p* < 0.05, \*\**p* < 0.01, \*\*\**p* < 0.001 represent the significance from a one-way ANOVA. **(E)** Western blot analysis showing the expression of S100A4 in tumor initiation and tumor progression. The densitometric analyses compare the protein expression relative to β-actin (n = 4). Mean ± SEM; \**p* < 0.05, \*\**p* < 0.01 represent the significance level from an unpaired *t*-test. **(F)** The expression of Vimentin and E-cadherin during tumor initiation and tumor progression is shown using real-time PCR in control and niclosamide-treated tumors (*n* = 4). Mean ± SEM; \**p* < 0.05, \*\*\**p* < 0.001, \*\*\*\**p* < 0.0001 represent the significance level from an unpaired *t*-test. Indicated (n) represents the independent experiments as biological replicates.

### 3. CAFs create an immunosuppressive tumor microenvironment by inhibiting NK cell activity

Previous studies have demonstrated that CAFs can suppress NK cell activity via inducing ferroptosis in gastric cancer^39^. Analysis of single-cell RNA sequencing datasets^40^ from multiple breast cancer patient tumor microenvironments revealed an inverse correlation (although not significant) between CAF abundance and NK cell populations (**Fig. 4A**). Extending these observations, we examined the immune landscape of human HER2-low breast tumors across different stages of disease progression and compared them with other breast cancer subtypes. Immunostaining of HER2-low human tumor specimens demonstrated that increased abundance of S100A4-enriched CAFs was inversely associated with NK cell populations (**Fig. 4B**). Single-cell Spatial transcriptomics with HER2-low human tumor samples showed that CAFs form dense, collagen-rich stromal barriers at the invasive front, which are associated with reduced infiltration of immune cells, including NK cells (**Fig. 4C**). Multiplex immunostaining of human HER2-low tumor samples revealed that HER2-low tumors exhibited an inverse association between S100A4-enriched CAF abundance and immune-cell infiltration. Specifically, tumors with higher S100A4-enriched CAF abundance showed reduced infiltration of antitumor immune populations, including T cells, M1 macrophages, and NK cells.

**Figure 4:**
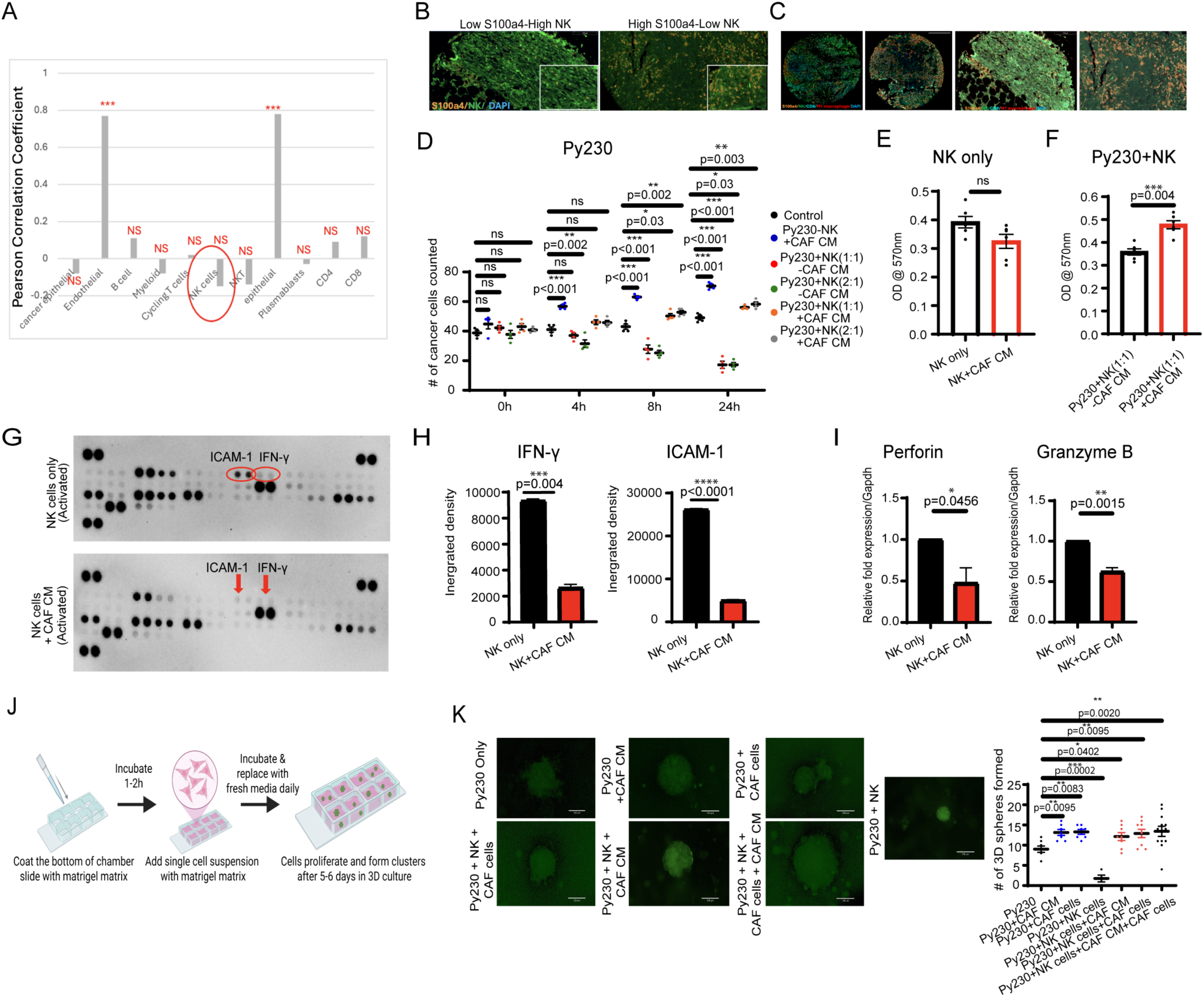
S100A4+ CAF CM inhibits cytotoxicity and the mechanisms of NK cells. **(A)** Analysis of scRNA-seq datasets from multiple breast cancer patient tumor microenvironments revealing an inverse correlation between CAF abundance and NK cell populations. Mean ± SEM; \*\*\**p* < 0.001 represents the significance level from a one-way ANOVA. **(B)** Multiplex immunostaining of HER2-low human tumor specimens inverse association with NK cell populations. Scale bar: 130µm. **(C)** Spatial mapping of CAFs’ association with reduced infiltration of immune cells. Scale bar: 170µm. **(D)** GFP-tagged Py230 cancer cells co-cultured with spleen NK cells at a 1:1 and 2:1 ratio in the presence and absence of S100A4-enriched CAF CM over 24 h. Mean ± SEM; \**p* < 0.05, \*\**p* < 0.01, \*\*\**p* < 0.001 represent the significance level from a two-way ANOVA. **(E)** Cell proliferation assay of NK cells only in the presence or absence of CAF CM and **(F)** addition of Py230 cancer cells. Mean ± SEM; \*\*\**p* < 0.001 represents the significance level from an unpaired *t*-test. **(G, H)** IL-12 and IL-15-activated spleen NK cells were incubated in the presence and absence of S100A4-enriched CAF CM for 24 h, and these samples were analyzed using a Proteome Profiler Mouse Cytokine Array Kit, Panel A. Mean ± SEM; \*\*\**p* < 0.001, \*\*\*\**p* < 0.0001 represent the significance level from an unpaired *t*-test. **(I)** The expression of Perforin and Granzyme B is shown using real-time PCR in IL-12 and IL-15-activated spleen NK cells, which were incubated in the presence and absence of S100A4-enriched CAF CM (*n* = 4). Mean ± SEM; \**p* < 0.05, \*\**p* < 0.01 represent the significance level from an unpaired *t*-test. **(J)** Schematic diagram of 3D cell culture. (**J**, Created with BioRender.com). **(K)** Images along with the number of 3D cell culture GFP-tagged Py230 co-cultured with spleen NK cells in the presence and absence of S100A4-enriched CAF CM and CAF cells at a 1:1 ratio (n=4). Mean ± SEM; \**p* < 0.05, \*\**p* < 0.01, \*\*\**p* < 0.001 represent the significance level from a one-way ANOVA. Scale bar: 120µm. Indicated (n) represents the independent experiments as biological replicates.

To directly assess the impact of S100A4-enriched CAFs on NK cell function, NK cells isolated^41^ from the spleen of wild-type C57BL/6J mice were treated with CAF-conditioned media (CAF-CM). CAF- CM markedly impaired NK cell–mediated cytotoxicity against Py230 tumor cells (**Fig. 4D**) and mammosphere formation over 24 h (**Supplementary Fig. 6A).** NK cell proliferation was decreased when CAF-CM was present, although not significantly (**Fig. 4E**). While activated NK cells effectively suppressed Py230 cancer cell proliferation, these properties were significantly attenuated in the presence of CAF-CM (**Fig. 4F**). Collectively, these findings indicate that S100A4-enriched CAFs suppress NK cell effector function. Mechanistically, NK cell cytotoxicity is mediated in part through interferon-γ (IFN-γ) secretion^42^. Cytokine–chemokine array analysis revealed that CAF-CM significantly reduced IFN-γ release (**Fig. 4G**), along with additional NK cell–associated effector molecules, including ICAM, from activated NK cells (**Fig. 4H**). Analysis of additional cytolytic effectors revealed that perforin and granzyme B expression in NK cells was significantly reduced following incubation with CM (**Fig. 4I**). Together, these findings demonstrate that S100A4-enriched CAFs suppress NK cell–mediated tumor clearance through a paracrine mechanism independent of direct cell–cell contact, establishing CAF-derived soluble factors as key mediators of immune evasion in HER2-low breast cancer. In a physiologically relevant 3D co-culture system (**Fig. 4J**) comprising Py230 tumor cells, primary NK cells, and S100A4-enriched CAFs, NK cells efficiently suppressed Py230 cell growth under monoculture conditions. Incorporation of S100A4-enriched CAFs (co-culture), S100A4-enriched CAF-conditioned media (paracrine), or both markedly attenuated NK cell– mediated cytotoxicity, resulting in enhanced cancer cell survival (**Fig. 4K**). This indicates that both contact-dependent and paracrine mechanisms cooperatively suppress NK-cell activity. Together, these findings demonstrate that S100A4-enriched CAFs suppress NK cell cytotoxicity through both soluble factor– mediated paracrine signaling and direct cell–cell interactions, thereby establishing a multifaceted immunosuppressive niche that promotes tumor cell survival in HER2-low breast cancer.

### 4. S100A4-enriched CAFs inhibit NK cell activity by altering amino acid metabolism in tumors

To identify the CAF-derived soluble factor(s) responsible for suppressing NK-cell activity, we next asked whether the inhibitory activity of CAF-CM was mediated by soluble proteins or by smaller molecular components, such as metabolites. CAF-CM was fractionated using molecular weight cutoff columns (3 kDa filter) to separate low-molecular-weight molecules from the protein-enriched fraction (**Fig. 5A**). The low-molecular-weight flow-through fraction (most proteins, cytokines, growth factors, and larger peptides will be retained in the 3 kDa fraction, while small metabolites should pass into the flow-through) retained NK-suppressive activity, supporting a metabolite-driven mechanism shown in the cell proliferation assay (**Fig. 5B).**

**Figure 5.**
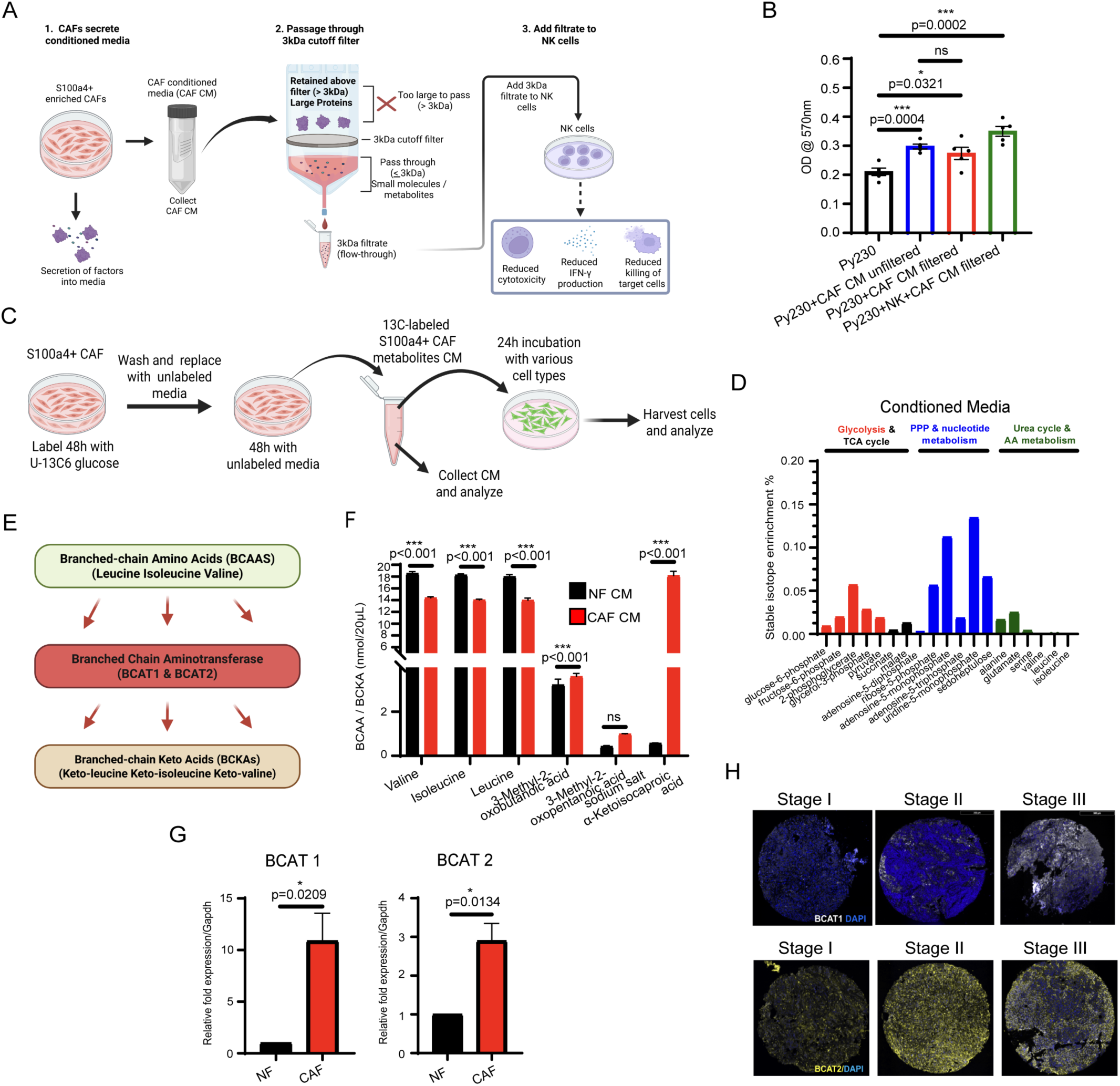
S100A4 CAFs’ effect on metabolic rewiring. **(A)** A schematic diagram of the CAF-CM fractionated using molecular weight cutoff columns (3 kDa filter) to separate low-molecular-weight molecules from the protein-enriched fraction, and **(B)** an MTT assay (n=5) analyzing cell proliferation using molecular weight cutoff columns (3 kDa filter) on CAF-CM. (**A**, Created with BioRender.com). Mean ± SEM; \**p* < 0.05, \*\*\**p* < 0.001 represent the significance level from a one-way ANOVA. **(C)** A schematic diagram of the TRACER experiment done by assessing carbon flux from ^13^C_6_-glucose into central energy metabolism pathways: Glycolysis, TCA Cycle, Pentose Phosphate Pathway, Amino Acids, and Urea. (24 h incubation) (n = 1 case). (**C**, Created with BioRender.com). **(D)** S100A4-enriched CAFs were fed ^13^C_6_-glucose tracer, and the conditioned media were collected for assessment. **(E)** Schematic diagram of branched-chain amino acids (BCAA) conversion to branched-chain keto acids (BCKA) by the enzyme branched-chain aminotransferase (BCAT1 & BCAT2). (**E**, Created with BioRender.com). **(F)** BCAA and BCKA amino acid profiling analysis of NF CM vs CAF CM (n=3). Mean ± SEM; \*\*\**p* < 0.001 represents the significance level from a one-way ANOVA. **(G)** The expression of BCAT1 and BCAT2 is shown using real-time PCR in S100A4-enriched CAF cells (n=4). Mean ± SEM; \**p* < 0.05 represents the significance level from an unpaired *t*-test. **(H)** Immunostaining of human BCAT 1&2 in the studied HER2-low tumors. Scale bar: 300µm. Indicated (n) represents the independent experiments as biological replicates.

An earlier study showed that CAFs deregulate glucose metabolism and facilitate TNBC progression^43^. Studies suggest a lactate shuttle between CAFs and cancer cells in the TME^44^. To define the metabolic exchange between CAFs and HER2-low cancer cells, we evaluated the fate of labeled metabolites in cancer cells, and NK cells fed with the metabolic products of CAFs (**Fig. 5C**). Uniformly labeled ^13^C glucose ([U-^13^C] glucose) was fed to CAFs isolated from 8-weeks old tumors for 48 h, and utilization of glucose by CAFs was evidenced by labeled metabolites implicated in glycolysis, the TCA cycle, PPP, nucleotide metabolism, aa metabolism, and urea cycle. Next, 48 h after exposure to ^13^C glucose, CAFs were washed and then cultured for an additional 48 h in unlabeled glucose medium. The CAF-secreted labeled metabolome (in CM) revealed metabolites involved in PPP, the TCA cycle, and nucleotide metabolism (**Fig. 5B**). The CM containing CAF-labeled metabolites was also used to feed Py230 cells and NK cells for 24 hrs. Py230 cells and NK cells were harvested, and the ^13^C-labeled metabolites derived from these cells were measured (**Supplementary Fig. 6B, C**). Analysis of metabolite patterns showed a decrease in the abundance of branched-chain amino acids (BCAAs) in the CAF-conditioned media **(Fig. 5D).** These experiments indicate that cancer cells and immune cells use CAF-derived metabolites to fuel their metabolic activity. Branched-chain amino acids (BCAAs; leucine, isoleucine, and valine) are essential amino acids that must be obtained from the diet^45,46^. Cancer cells exhibit increased uptake of BCAAs, and the branched-chain aminotransferases (BCATs), which catalyze the first step in BCAA catabolism, are frequently overexpressed in tumors^47–49^. The enzymes Branched-Chain Amino Acid Transaminase 1 (BCAT1) and Branched-Chain Amino Acid Transaminase 2 (BCAT2) convert BCAAs into branched-chain α-ketoacids (BCKAs) through a reversible transamination reaction^50^ (**Fig. 5E**). Notably, a recent study demonstrated markedly elevated BCAA catabolic flux in CAFs in pancreatic cancer, with stromal BCKA production critically dependent on BCAT1-mediated transamination^51^. While tumor-derived metabolites are known to suppress NK cell function^52^, the role of BCKAs as stromal-derived immunosuppressive metabolites remains unexplored. Whether CAF-derived BCKAs directly impair NK cell cytotoxicity represents a critical and unaddressed gap in the field. Therefore, we further measured the abundance of BCAAs and BCKAs in the CM of the S100A4-enriched CAFs and CM from NFs via targeted metabolomics using Mass spectrometry (**Supplementary Fig. 6D**). Our study showed that CM from S100A4-enriched CAFs produced significantly lower BCAAs compared to the control CM (**Fig. 5F**). Further HPLC analysis showed a significant decreased in BCAA and an increased BCKA (especially α-ketoisocaproic acid) in CAF-CM compared to control CM (**Fig. 5F**). Because NK cell cytotoxicity is mediated in part by IFN-γ secretion, we next examined whether BCAA or BCKAs, directly inhibit NK cells cytotoxic properties by affecting IFN-γ production. ELISA analysis revealed that BCAA treatment did not alter IFN-γ secretion from activated NK cells, whereas BCKA treatment significantly suppressed IFN-γ production (**Supplementary Fig. 7A**). Real-time PCR study showed a significant increase in both BCAT1 and BCAT2 transcript levels in the S100A4-enriched CAF population compared to NFs (**Fig. 5G**). Additionally, western blot analysis showed a significant increase in both BCAT1 and BCAT2 downstream targets, FOXM1 and SREBP1, in CAF samples compared to the BCAT1-and BCAT2-inhibited samples (**Supplementary Fig. 7B**). Immunostaining of human HER2-low breast tumor samples revealed elevated expression of BCAT1 and BCAT2 in advanced-stage tumors (**Fig. 5H**).

### 5. Targeting BCAT1/2 to inhibit CAF-induced HER2-low mammary tumorigenesis

A recent study demonstrated that CAFs in TNBC suppress NK cell activity through the NKG2D–DNAM-1 axis^53^. Here, we identify a distinct metabolic mechanism by which S100A4-enriched CAFs inhibit NK cell cytotoxic function through modulation of amino acid metabolism. Because S100A4-enriched CAFs upregulate both BCAT1 and BCAT2, we independently inhibited each enzyme and assessed the impact on NK cell cytotoxicity following culture with CM from control or treated CAFs. Pharmacologic inhibition of BCAT2 in S100A4-enriched CAFs using BCATc Inhibitor 2^54^ did not significantly alter NK cell–mediated cytotoxicity to the control (**Fig. 6A**). In contrast, inhibition of BCAT1 using Erg240^55^ markedly restored NK cell cytotoxic function in mammosphere formation, as CM from Erg240-treated S100A4-enriched CAFs failed to suppress NK cell–mediated tumor cell killing (**Fig. 6B**). Consistent with these *in vitro* findings, *in vivo* treatment of Py230 tumor–bearing mice at 3 weeks post-implantation with Erg240 resulted in a significant reduction in tumor volume, suggesting restoration of NK cell antitumor activity through BCAT1 inhibition (**Fig. 6C**). Importantly, NK cells cultured with conditioned media from Erg240-treated CAFs, which exhibited reduced BCKA levels (**Fig. 6D**), retained their ability to secrete IFN-γ (**Fig. 6E**). These findings indicate that BCAT1 inhibition prevents CAF-mediated suppression of NK-cell cytotoxicity by blocking BCAA-to-BCKA conversion. To determine whether the antitumor effects of BCAT1 inhibition are mediated through NK cells, Py230 HER2-low mammary tumor cells were orthotopically implanted into the mammary fat pads of immunocompetent C57BL/6J female mice. Once tumors became palpable, mice were randomized into four groups receiving vehicle or the BCAT1 inhibitor Erg240, each in combination with either isotype control antibody or NK-cell depletion antibody. NK cells were depleted using anti-NK1.1 administered beginning one day prior to treatment initiation and continued at regular intervals throughout the study. BCAT1 inhibition was found to suppress tumor growth in NK-intact mice, whereas NK cell depletion diminished this therapeutic effect, thereby demonstrating that the antitumor activity of BCAT1 inhibition is mediated, at least in part, through restoration of NK-cell function (**Fig. 6F, G**).

**Figure 6:**
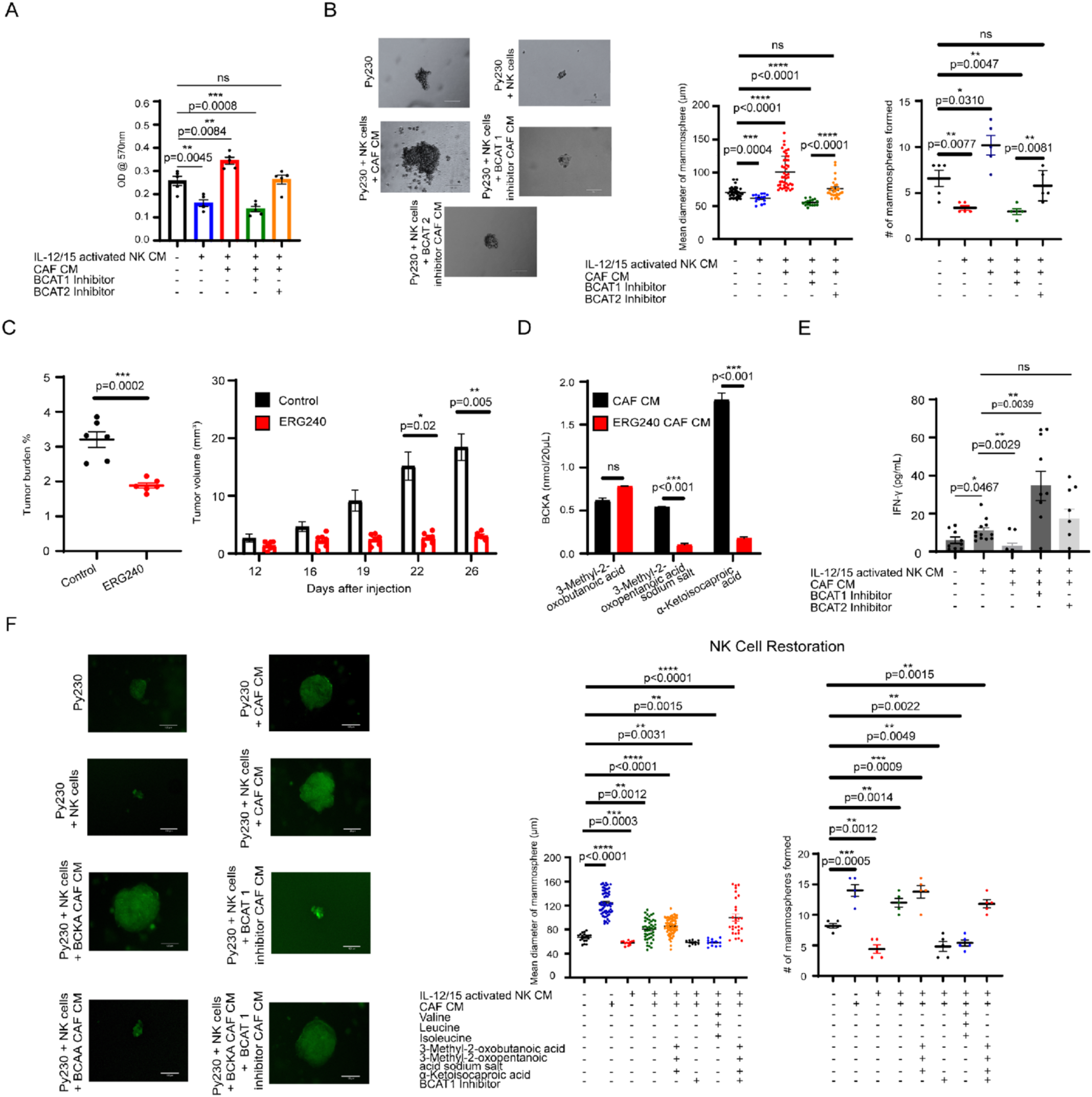
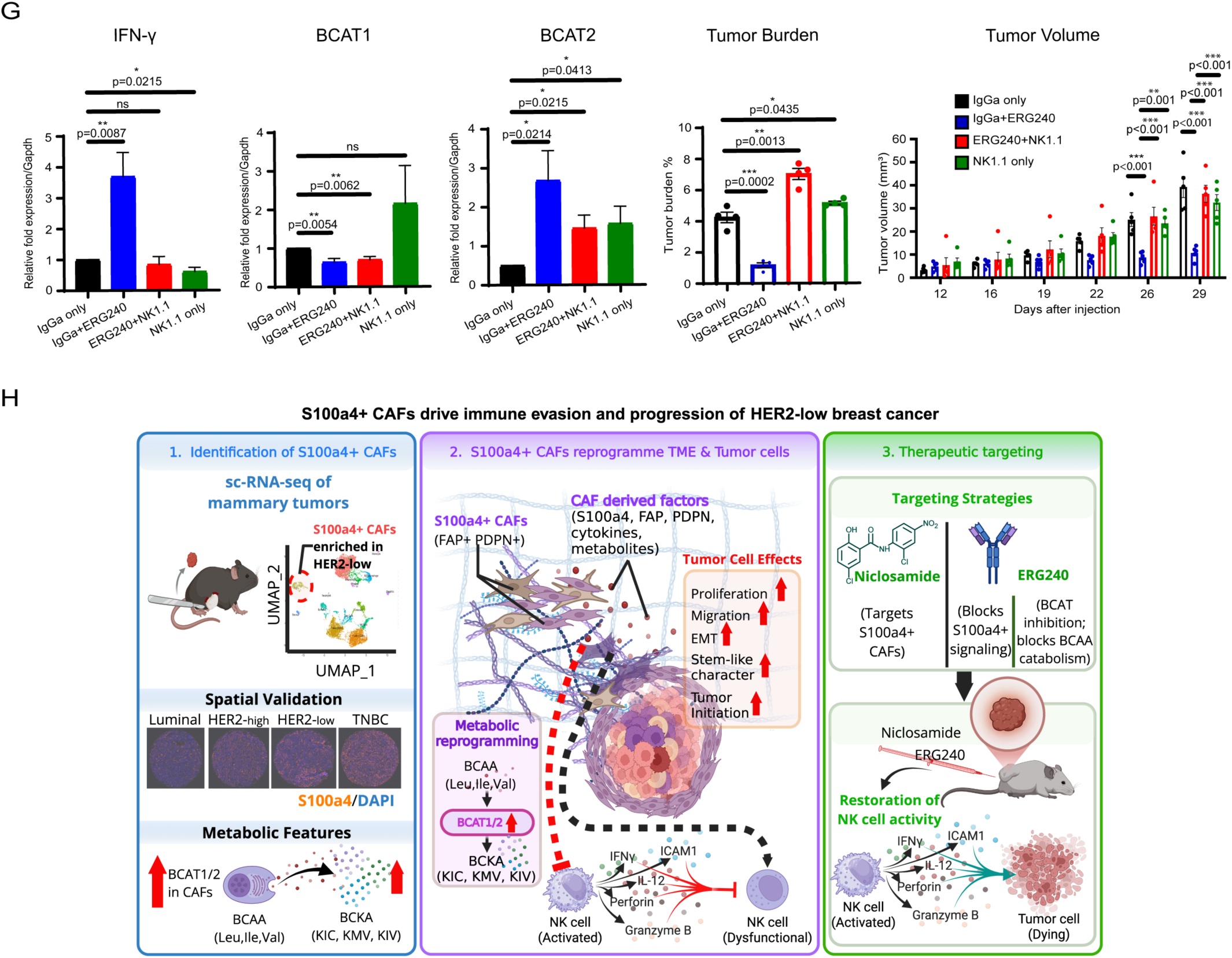
NK cell restoration via inhibition of the enzyme BCAT1. **(A)** MTT assay analyzing the cell proliferation of Py230 cancer cells in the presence or absence of IL-12 and IL-15-activated NK cells, CAF CM, and BCAT1 & 2 inhibitors (n=5). Mean ± SEM; \*\**p* < 0.01, \*\*\**p* < 0.001 represent the significance level from a one-way ANOVA. **(B)** Images along with mean diameter and number of mammospheres of Py230 incubated in the presence and absence of IL-12 and IL-15-activated spleen NK cells and in the presence and absence of S100A4-enriched CAF CM with BCAT1 inhibitor and BCAT2 inhibitor (n=5). Mean ± SEM; \**p* < 0.05, \*\**p* < 0.01, \*\*\**p* < 0.001, \*\*\*\**p* < 0.0001 represent the significance level from a one-way ANOVA. Scale bar: 120µm. **(C)** HER2-low tumor data from i.p. injection of C57BL/6J mice with ERG240 in tumor burden percentage (tumor-to-body-weight ratio) and tumor volume (n=7-12 for each experimental condition). Tumor burden: Mean ± SEM; ***p < 0.001 represents the significance level from an unpaired two-sided *t*-test. Tumor volume: Mean ± SEM; *p < 0.05, \*\**p* < 0.01 represent the significance level from a one-way ANOVA. **(D)** BCKA amino acid profiling analysis of CAF CM vs ERG240-enriched CAF CM (n=3). Mean ± SEM; \**p* < 0.05 represents the significance level from a one-way ANOVA. **(E)** ELISA MAX™ Deluxe Set Mouse IFN-γ was utilized to measure IFN-γ levels in cell culture supernatants of IL-12 and IL-15-activated spleen NK cells under various incubation conditions, utilizing BCAT 1 & 2 inhibitors (*n* = 9-12). \**p* < 0.05, \*\**p* < 0.01 represent the significance level from a one-way ANOVA. **(F)** Images along with mean diameter and number of mammospheres of GFP-tagged Py230 incubated in the presence and absence of IL-12 and IL-15-activated spleen NK cells and in the presence and absence of S100A4-enriched CAF CM with BCAAs, BCKAs, and BCAT1 inhibitor (n=5). \*\**p* < 0.01, \*\*\**p* < 0.001, \*\*\*\**p* < 0.0001 represents the significance level from a one-way ANOVA. Scale bar: 120µm. **(G)** The expression of IFN-γ, BCAT1, and BCAT2 is shown using real-time PCR in IgGa control, ERG240, and NK1.1 treatments (n=5). Mean ± SEM; *p < 0.05, **p < 0.01, ***p < 0.001, ****p < 0.0001 represent the significance level from a one-way ANOVA. **(G)** Tumor burden percentage (tumor-to-body-weight ratio) and tumor volume (n=5 for each experimental condition) for IgGa control, ERG240, and NK1.1 treatment via i.p. injection. Tumor burden: Mean ± SEM; *p < 0.05, **p < 0.01, ***p < 0.001 represent the significance level from an unpaired two-sided *t*-test. Tumor volume: Mean ± SEM; \*\**p* < 0.01, \*\*\**p* < 0.001 represent the significance level from a one-way ANOVA. **(H)** Schematic diagram of this study identifying a previously unrecognized CAF-driven metabolic mechanism of innate immune suppression in HER2-low breast cancer. (Created with BioRender.com).

Collectively, these results identify S100A4-enriched CAFs as a tumor-promoting stromal population in HER2-low breast cancer and define BCAT1-dependent BCKA production as a metabolic mechanism of NK-cell suppression. By linking stromal CAF expansion in human HER2-low tumors to impaired NK-cell immunity and demonstrating that BCAT1 inhibition restores antitumor activity *in vivo*, our findings reveal a targetable CAF-driven metabolic immune checkpoint in HER2-low breast cancer.

## Discussion

HER2-low breast cancer has emerged as a clinically important disease category; however, the stromal and immune mechanisms that shape its progression remain incompletely understood. In this study, we identify a CAF-driven metabolic–immune axis (**Fig. 6H**) that suppresses NK-cell–mediated tumor surveillance in HER2-low breast cancer. Using human tumor profiling, murine HER2-low tumor models, functional co-culture systems, metabolomics, and pharmacologic intervention, we show that S100A4-enriched CAFs expand during HER2-low tumor progression and establish an immunosuppressive tumor microenvironment. Mechanistically, these CAFs upregulate BCAT1-dependent branched-chain amino acid metabolism, resulting in accumulation of branched-chain keto acids that impair NK-cell IFN-γ production and cytotoxicity. These findings define a stromal metabolic checkpoint through which CAFs suppress innate antitumor immunity and promote HER2-low tumor progression.

A major finding of this study is the identification of S100A4-enriched CAFs as a dominant stromal population associated with HER2-low tumor advancement. Single-cell spatial transcriptomic analysis of human HER2-low tumors revealed progressive expansion of the CAF compartment, which was associated with reduced immune-cell infiltration and a more immunosuppressive tumor microenvironment. Multiplex immunostaining further showed that S100A4, but not PDPN or FAP, increased progressively from stage I to stage III HER2-low tumors, indicating that tumor progression is accompanied by selective enrichment of a specific CAF state rather than uniform expansion of all CAF populations. This finding is important because CAFs are highly heterogeneous^56^, and bulk stromal abundance alone does not explain their functional contribution to tumor biology. Our data suggest that S100A4-enriched CAFs represent a clinically relevant stromal subset linked to immune exclusion and aggressive disease behavior in HER2-low breast cancer.

The inverse relationship between S100A4-enriched CAF abundance and immune-cell infiltration provides an important human-tumor context for the functional studies. Regions enriched for S100A4⁺ CAFs showed reduced infiltration of NK cells, T cells, and macrophages, suggesting that these CAFs contribute to the formation of immune-restricted stromal niches. While CAFs have been widely implicated in T-cell exclusion and therapy resistance^57–59^, their role in regulating NK-cell immunity in HER2-low breast cancer has remained poorly defined. NK cells are central mediators of innate tumor surveillance and are particularly relevant in settings where tumor antigen presentation may be impaired^60,61^. Our data show that S100A4-enriched CAFs suppress NK-cell cytotoxicity through both soluble factor–mediated and contact-dependent mechanisms. CAF-conditioned media was sufficient to impair NK-cell killing, reduce IFN-γ secretion, and decrease perforin and granzyme B expression, indicating that CAF-derived paracrine factors directly compromise NK-cell effector function.

Although CAF-mediated suppression of immune cells has been reported, most studies have focused on contact-dependent mechanisms or cytokine signaling pathways^62^. Our work extends these findings by demonstrating that CAF-derived metabolites are sufficient to suppress NK-cell function, independent of direct cell–cell interactions. Using conditioned media and co-culture systems, we show that CAFs suppress NK-cell cytotoxicity through both paracrine and contact-dependent mechanisms, with soluble factors playing a dominant role. These findings position CAFs as active regulators of the metabolic microenvironment that shapes immune cell function. An important mechanistic advance of this study is the identification of altered branched-chain amino acid metabolism as a mediator of CAF-induced NK-cell suppression. Metabolic tracing and targeted metabolomics showed that S100A4-enriched CAFs altered BCAA metabolism and generated increased levels of BCKAs. Although BCAA metabolism has been implicated in tumor growth and cancer-cell fitness^63,64^, our findings extend this concept by demonstrating that stromal BCAA catabolism can suppress antitumor immunity. Notably, BCAA treatment alone did not impair NK-cell IFN-γ secretion, whereas BCKA treatment significantly suppressed IFN-γ production. This distinction suggests that the immunosuppressive activity is not caused simply by altered amino acid availability, but by CAF-dependent metabolic conversion of BCAAs into immunomodulatory keto-acid products.

Our data further identify BCAT1 as a critical enzymatic regulator of this stromal metabolic immune checkpoint. S100A4-enriched CAFs expressed increased BCAT1 and BCAT2, but pharmacologic inhibition of BCAT1, rather than BCAT2, restored NK-cell cytotoxic function. Conditioned media from BCAT1-inhibited CAFs failed to suppress NK-cell tumor killing and preserved IFN-γ secretion, supporting a model in which BCAT1-dependent BCKA production is required for CAF-mediated NK-cell dysfunction. *In vivo*, BCAT1 inhibition suppressed HER2-low tumor growth, and this antitumor effect was diminished by NK-cell depletion, demonstrating that restoration of NK-cell activity contributes directly to the therapeutic effect. Together, these results establish a causal link between CAF metabolic programming, NK-cell suppression, and HER2-low tumor progression.

These findings have several translational implications. First, they suggest that S100A4-enriched CAFs may serve as a stromal biomarker of immune exclusion in HER2-low tumors. Second, they identify BCAT1-dependent BCAA metabolism as a potentially targetable pathway within the tumor stroma. Third, they raise the possibility that inhibiting CAF metabolic suppression could improve innate immune surveillance and enhance responses to therapies that depend on immune engagement. This may be particularly relevant for HER2-low breast cancer, where antibody–drug conjugates have improved clinical outcomes, but resistance remains a major challenge. Because NK cells can contribute to antibody-dependent cellular cytotoxicity and broader innate immune control, relieving CAF-mediated NK-cell suppression may provide a rational strategy to improve antitumor immunity in HER2-low disease.

This study also highlights the importance of functionally defining CAF subsets rather than treating CAFs as a single stromal population. Inhibiting FAP⁺ or PDPN⁺ CAF populations did not significantly reduce the tumor-promoting activity of CAF-conditioned media, whereas pharmacologic inhibition or siRNA-mediated suppression of S100A4 attenuated CAF-induced aggressive behavior in HER2-low cancer cells. These findings support the concept that CAF-directed therapies must be subset-specific and mechanism-based. Broad depletion of fibroblasts may produce unpredictable or even detrimental effects, whereas targeting a defined tumor-promoting CAF program, such as the S100A4–BCAT1–BCKA axis, may offer a more precise therapeutic opportunity.

There are limitations to this study. Although our human tumor data demonstrate an association between S100A4⁺ CAF expansion, BCAT1 expression, and reduced immune infiltration, larger patient cohorts will be required to determine whether this axis predicts clinical outcome or therapeutic resistance in HER2-low breast cancer. Future studies should also define whether stromal BCAT1 expression correlates with response to antibody–drug conjugates, chemotherapy, or immunotherapy. In addition, while our data support BCKAs as suppressors of NK-cell function, the downstream signaling pathways by which BCKAs impair IFN-γ production and cytolytic activity remain to be fully defined. Finally, determining whether this CAF metabolic program operates in other breast cancer subtypes or additional solid tumors will be important for establishing the broader relevance of this mechanism.

In summary, our study identifies S100A4-enriched CAFs as a functionally important stromal population that expands during HER2-low breast tumor progression and promotes immune exclusion. We demonstrate that these CAFs suppress NK-cell antitumor activity through BCAT1-dependent conversion of BCAAs into BCKAs, thereby establishing a metabolically immunosuppressive niche. Pharmacologic inhibition of BCAT1 restores NK-cell function and suppresses tumor growth in an NK-cell–dependent manner. These findings reveal a CAF-driven metabolic immune checkpoint in HER2-low breast cancer and nominate the S100A4–BCAT1–BCKA axis as a targetable stromal vulnerability to restore innate antitumor immunity.

## Supporting information

Carter Supplementary doc

## Materials availability

This study did not generate any new materials.

## Data availability

The Xenium data generated in this study will be made publicly available upon acceptance of the manuscript.

## Code availability

The code will be available upon acceptance of this paper.

## Acknowledgements

This work was funded by the Texas A&M University Faculty Development Grant (to T.R.S.), PRISE grant (to T.R.S.), and Strategic Transformative Research Funds (STRP to T.R.S.), and 1R01GM163238 (to T.R.S.). This work was supported, in part, by a Pilot grant from NIEHS P30ES029067 (to T.R.S.). We would like to thank Omics Empower Inc. for helping with the bioinformatics analyses. We would like to thank Dr. Richard Gomer and Dr. Darrell Pilling for their invaluable suggestions and insightful feedback.

## Author contributions

Conceptualization: T.R.S, and K.C. Methodology: K. C., O.O, M.S., S.A., Investigation: K. C., O.O, M.S., S.A., C.N., M.N., A.A, D.F., G. N., C. K. Data analysis: K.C., M.S., J.C., Supervision: T.R.S., K. C. Writing-original draft: T.R.S., and, K.C, All authors reviewed, read, and agreed to publish the manuscript.

## Methods

### Mice

C57BL/6J (Jax # **000664)**. Mice were sacrificed by cervical dislocation for tumor or normal tissue harvesting. All animal care and treatments were in accordance with the Texas A&M University Animal Care and Use Committee under protocol #2022-0094.

### Cell lines and media

Py230, BT-20, 4T1, MDA-MB-231, MCF-7, and SK-BR-3 cell lines were purchased from the ATCC. Py230, a mouse HER2-Low cell line, was grown in F12K Medium (ATCC; Manassas, VA, USA) supplemented with 5% Fetal Bovine Serum (FBS) (ATCC; Manassas, VA, USA), 0.1% MITO+ Serum Extender (Corning #355006). Py230 cells do not maintain their properties without the MITO+ Serum Extender. BT-20, a human HER2-Low cell line, was grown in Eagle’s Minimum Essential Medium (EMEM) (ATCC; Manassas, VA, USA) supplemented with 10% Fetal Bovine Serum (FBS) (ATCC; Manassas, VA, USA), 1% penicillin– streptomycin (Gibco™, Thermo Fisher Scientific, Inc., Waltham, MA, USA). 4T1, a mouse triple-negative cell line, was grown in Roswell Park Memorial Institute Medium (RPMI 1640) (ATCC; Manassas, VA, USA) supplemented with 10% Fetal Bovine Serum (FBS) (ATCC; Manassas, VA, USA), 1% penicillin– streptomycin (Gibco™; Thermo Fisher Scientific, Inc., Waltham, MA, USA). MDA-MB-231, a human triple-negative cell line, was grown in Dulbecco’s Modified Eagle’s Medium (DMEM) (ATCC; Manassas, VA, USA) supplemented with 10% Fetal Bovine Serum (FBS) (ATCC; Manassas, VA, USA), 1% penicillin– streptomycin (Gibco™; Thermo Fisher Scientific, Inc., Waltham, MA, USA). MCF-7, an ER+ cell line, was grown in Dulbecco’s Modified Eagle’s Medium (DMEM) (ATCC; Manassas, VA, USA) supplemented with 10% Fetal Bovine Serum (FBS) (ATCC; Manassas, VA, USA), 1% penicillin–streptomycin (Gibco™; Thermo Fisher Scientific, Inc., Waltham, MA, USA). SK-BR-3, an HER2+ cell line, was grown in McCoy’s 5A (ATCC; Manassas, VA, USA) supplemented with 10% Fetal Bovine Serum (FBS) (ATCC; Manassas, VA, USA), 1% penicillin–streptomycin (Gibco™; Thermo Fisher Scientific, Inc., Waltham, MA, USA).

### Flow Activated Cell Sorting

Normal mammary fibroblasts (NF) were collected from C57BL/6J females. Fat pad tissue was minced and dissociated using the gentleMACS dissociator with an enzymatic digestion solution consisting of 5 mg Collagenase Type IV, Cls IV (MilliporeSigma, #C4-28; Burlington, MA), 25 mg Dispase II (MilliporeSigma, #D4693-1G; Burlington, MA), 20 uL DNase I recombinant (MilliporeSigma, #04536282001; Burlington, MA) in 5 mL of RPMI 1640. The tissue was incubated (37 °C, 5% CO_2_) for 45 minutes to 1 hour while spinning. The samples were filtered through a cell strainer into 10 mL of RPMI 1640. The cell suspension was pelleted, centrifuged at 300 × g for 7 minutes at 4°C, followed by Red Blood Cell lysis treatment (BioLegend, 420302). 10 mL of chilled wash buffer (4mg of BSA in 10 mL of PBS) was added and centrifuged at 300 × g for 10 minutes at 4°C. The supernatant was removed, and 5 mL of 5 mL of wash buffer was added to count cells, followed by staining with FITC anti-mouse CD326, Alexa Fluor 594 anti-CD45, and Pacific Blue anti-CD31 antibodies for exclusion. Finally, FACS was used to collect NF cells. Following antibody incubation, the cells were washed with FACS buffer (100µL of FBS in 10mL of PBS). DAPI or 7-AAD was added before FACS sorting to distinguish between live and dead cells. CAFs were collected from C57BL/6J female mice. Tumors were minced and dissociated using the gentleMACS dissociator^65^ with an enzymatic digestion solution consisting of 5 mg Collagenase Type IV, Cls IV (MilliporeSigma, #C4-28; Burlington, MA), 25 mg Dispase II (MilliporeSigma, #D4693-1G; Burlington, MA), 20 uL DNase I recombinant (MilliporeSigma, #04536282001; Burlington, MA) in 5 mL of RPMI 1640. The tissue was incubated (37 °C, 5% CO_2_) for 45 minutes to 1 hour while spinning. The samples were filtered through a cell strainer into 10 mL of RPMI 1640. The cell suspension was pelleted, centrifuged at 300 × g for 7 minutes at 4°C, followed by Red Blood Cell lysis treatment (BioLegend, 420302). 10 mL of chilled wash buffer (4mg of BSA in 10 mL of PBS) was added and centrifuged at 300 × g for 10 minutes at 4°C. The supernatant was removed, and 5 mL of wash buffer was added to count cells, followed by staining: FITC anti-mouse CD326 (BioLegend, #118207, San Diego, CA, USA), Alexa Fluor 594 anti-CD45 (BioLegend, #103144, San Diego, CA, USA), Pacific Blue anti-CD31(BioLegend, #102421, San Diego, CA, USA), Fibroblast Activation Protein alpha/FAP Antibody (983802) [Alexa Fluor® 647] (Novus Biologicals, #FAB9727R-100UG, Centennial, CO, USA), and APC/Fire™ 750 anti-mouse Podoplanin Antibody (Clone 8.1.1) (BioLegend, #127425, San Diego, CA, USA), and NG2/MCSP Antibody (546930) [Alexa Fluor® 647] (Novus Biologicals, # FAB6689R-100UG, Centennial, CO, USA) antibodies for exclusion. Finally, the CAF population was collected. Following antibody incubation, the cells were washed with FACS buffer (100µL of FBS in 10mL of PBS). DAPI or 7-AAD was added before FACS sorting to distinguish between live and dead cells.

### Conditioned Media Collection

Following initial isolation, once NFs or CAFs achieved 70-90% confluency in DMEM, the conditioned medium was collected, passed through a 0.45 µm filter, and centrifuged for 5 mins at 300g before use on various cell types or frozen at −80°C.

### 3 kDa MWCO filtering

Following initial isolation, once CAFs achieved 70-90% confluency in DMEM, the conditioned medium was collected and passed through an Amicon® Ultra Centrifugal Filter, 3 kDa MWCO (MilliporeSigma, # UFC900308; Burlington, MA), and centrifuged for 1 h at 4,000 g before use on various cell types or frozen at −80°C.

### S100A4 siRNA

S100A4 Mouse Pre-designed siRNA Set A (MedChemExpress, #HY-RS16679, Monmouth Junction, NJ, USA) was utilized to knock down S100A4 protein in culturing CAF cells. Following 24 h of the addition of Growth Medium (2 μL), 20 μM siRNA (4 μL), siRNA/mRNA Transfection Reagent (8 μL), and Reduced-serum medium (400 μL) to CAFs in a 6-well plate, the conditioned medium was collected, passed through a 0.45 µm filter, and centrifuged for 5 mins at 300g before use for downstream experiments or frozen at −80°C.

### NK Cell Isolation and Culture

Murine NK cells were isolated from the spleen of C57BL/6 mice using magnetic separation (Miltenyi Biotec, mouse NK Cell Isolation Kit, 130-115-818). Murine NK cells were grown in Roswell Park Memorial Institute Medium (RPMI 1640) (ATCC; Manassas, VA, USA) supplemented with 10% Fetal Bovine Serum (FBS) (ATCC; Manassas, VA, USA), 1% penicillin–streptomycin (Gibco™; Thermo Fisher Scientific, Inc., Waltham, MA, USA).

### Single-cell RNA sequencing (scRNA seq)

After brief washing with chilled sterile PBS, tumors were placed on ice in DMEM (ATCC, 30-2002) supplemented with 10% FBS (Sigma-Aldrich, F2442) and 1% antibiotic solution (Sigma-Aldrich, P4333) until further processing. Tumors were minced and mechanically dissociated using the gentleMACS Dissociator (Miltenyi Biotec, 130-093-235), followed by enzymatic digestion with the MACS Miltenyi Tumor Dissociation Kit for mice (Miltenyi Biotec, 130-096-730) according to the manufacturer’s protocol. The resulting single-cell suspension was transferred to the Texas A&M Institute for Genome Sciences and Society (TIGSS) for single-cell RNA sequencing. Single-cell sequencing libraries were prepared using the Chromium platform (10x Genomics) with the Single Cell 5′ v2 Full Kit (PN-1000263). Cell-type annotation was performed manually based on unique marker genes identified for each cluster following UMAP dimensionality reduction. Marker genes were cross-referenced with published literature on mouse mammary gland and mammary tumor tissues. Differentially expressed genes were identified using the MAST^66^ R package.

### Scratch Assay

The scratch assay was performed^31^ by seeding at a density of 0.01 × 10^6^ cells/well in 96-well plates and allowing 24 h to adhere in complete medium. Once a confluent monolayer was formed, a scratch was made using a sterile pipette tip, and the dish was treated with fresh medium. The medium was replaced at a 1:1 ratio of treatment (NF CM or CAF CM) with the normal culture medium, along with the control sample. The wound closure experiment was monitored and recorded for the next 31 h.

### MTT Assay

The cytotoxicity of all cell lines was analyzed by MTT cell proliferation assay^67^ kit (Cayman Chemical Company, #10009365; Ann Arbor, MI, USA) using the manufacturer’s standard protocol. The cells were seeded at a density of 0.03 × 10^6^ cells/well in 96-well plates (Thermo Fisher Scientific; Waltham, MA, USA) and were incubated overnight in a 5% CO_2_ atmosphere at 37 °C to allow attachment to the plate. The medium was replaced the following day at a 1:1 ratio of treatment (NF CM or CAF CM) with the normal culture medium, along with the control sample. Following 24 h of incubation with treatments, 10 μL of MTT reagent was added to each well and subjected to incubation for 4 h. After confirming the formation of the purple-colored formazan crystals under a microscope, the crystals were dissolved with 100 μL of crystal-dissolving solution (Cayman Chemical Company, #10009365; Ann Arbor, MI, USA). The plate was shaken on an orbital shaker for 5 mins before incubating for 30 minutes in a 5% CO_2_ atmosphere at 37 °C and recording the absorbance at 570nm using a spectrophotometric plate reader (Molecular Devices; San Jose, CA, USA).

### Mammosphere Assay

This assay was used to identify anchorage-independent growth of cancer stem cells^68^. The cells were seeded at a density of 0.0005 × 10^6^ cells/well in Nunclon™ Sphera™ 96-Well, Nunclon Sphera-Treated, U-Shaped-Bottom Microplates (Thermo Fisher Scientific; Waltham, MA, USA) supplemented at a 1:1 ratio with treatment of NF CM or CAF CM with the complete mammosphere media, along with the control sample. The plate was incubated for 5-7 days in a 5% CO_2_ atmosphere at 37°C to allow the anchorage-independent growth of cancer stem cells. After 5-7 days in culture, mammosphere images were captured by an Echo Rebel microscope (Echo, San Diego, CA, USA).

### Immunofluorescence Assay

The cells (3.5 × 10^5^ cells/mL) were seeded on a coverslip in a 6-well plate and incubated for 24 h at 37 °C with 5% CO_2_. Then, fixed with 4% paraformaldehyde (Santacruz Biotechnology Inc.; Dallas, TX, USA) at room temperature for 20min. The primary antibodies used for the immunofluorescence assay was FSP1/S100A4 Rabbit pAb (A1631f) (ABclonal, #A1631, 1:2000; Woburn, MA, USA), Fibroblast Activation Protein alpha/FAP Antibody (983802) [Unconjugated] (Novus Biologicals, #MAB9727-SP, 1:5000, Centennial, CO, USA), and Mouse Podoplanin Antibody (R&D Systems, # AF3244-SP, 1:5000; Minneapolis, MN, USA). The secondary antibody was anti-rabbit IgG (Alexa Fluor® 488) (Cell Signaling Technology, #4412S, 1:2000; Danvers, MA, USA); Goat Anti-Rat IgG H&L (Alexa Fluor® 594) (Abcam, # ab150160, 1:2000); and anti-mouse (Alexa Fluor® 647) (Cell Signaling Technology, #5059S, 1:2000; Danvers, MA, USA). The nuclei were visualized by DAPI (Prolong Gold Antifade with DAPI, Molecular Probes, #8961S; Thermo Fisher Scientific, Waltham, MA, USA), and the images were captured by the Echo Rebel microscope (Echo, San Diego, CA, USA).

### Multiplexing immunostaining on TMAs

Antibodies: Two separate panels were developed and optimized for use in this study. The following primary antibodies were utilized for both immunohistochemical and immunofluorescence staining: rabbit anti-human FoxP3 [BLR034F], rabbit anti-human CD163 [BLR087G], mouse anti-human PDPN [LpMab-12], rabbit anti-human FAP [BLR150J], mouse anti-human BCAT1 [OTI3F5], rabbit anti-human CD56 [BLR152J], rabbit anti-human BCAT2 [D8K30], rabbit anti-human S100A4 [D9F9D], rabbit anti-human CD86 [E2G8P], and mouse anti-human CD8a [144B]. Optimization: Optimization of antibody concentration and staining order was achieved using FFPE human immune tissue microarray serial sections. Each target was evaluated for overall signal: noise ratio, loss of signal intensity, elution efficiency, and overall autofluorescence via heat-induced epitope retrieval (HIER) methods. The final optimized panels were then subjected to the Spatomics’ CFP™ method of multiplex immunofluorescence (see below). Multiplex immunofluorescence staining was performed with Spatomics CFP™ cleavable dyes. This technology allows for numerous protein targets to be stained using CFPs, which generate strong, localized fluorescence through an HRP-catalyzed reaction between the dye and nearby residues. After imaging, the fluorescent tags are efficiently cleaved, and HRP is deactivated, allowing repeated cycles without compromising tissue quality or antigenicity. FFPE microarrays were baked for 60 minutes, deparaffinized in xylene, and rehydrated by serial passage through graded concentrations of ethanol. Endogenous peroxidase in tissues was blocked with 0.9% H_2_O_2_/methanol for 40 min. An initial HIER treatment was performed for 20 min at 92-96C in Tris-EDTA pH 9 buffer. Following HIER, slides were rinsed with DI water and cooled at RT for 20 minutes. Slides were blocked with 20% normal goat serum for 30 minutes before loading onto the Parhelia Spatial Station™ auto stainer. Primary antibodies were incubated for 20 min. Then, slides were rinsed with TBS for 10 min and incubated with HRP-conjugated secondary (A120-501P) for 20 min, followed by another 10 min rinse in TBS. Incubation with a CFP dye (670, 595, 555, 490, or 750) was done for two 10-min exchanges, followed by a 10-min rinse in DI water. After the first panel was stained, the slide was removed and incubated with DAPI for 10 minutes. The slide was mounted in VECTASHIELD Vibrance Antifade Mounting Medium (ThermoFisher Scientific) and imaged using the PhenoImager HT. Following imaging, the coverslip was removed and the first panel of bound primary and secondary antibodies was then cleaved off the tissue. The staining and imaging process was repeated for the second panel, resulting in two images [Panel 1 and Panel 2] of the same slide. Imaging: Akoya Biosciences’ PhenoImager HT (formerly known as Vectra Polaris Automated Quantitative Pathology Imaging System) was used for multispectral imaging at 40× magnification. Thereafter, whole slide images were uploaded for viewing on Pathcore.

### Spatial transcriptomics using Xenium platform

Human HER2-Low breast tumor formalin-fixed paraffin-embedded (FFPE) tissue sections were profiled using the Xenium Prime 5K Gene Expression assay with cell segmentation on the Xenium Analyzer (10x Genomics, Pleasanton, CA, USA). Samples were processed using Xenium slides, Xenium Prime Sample Prep Reagents (PN-1000720), Xenium Prime Cassettes and Inserts (PN-1000723), the Xenium Prime 5K Human Pan Tissue and Pathways Panel (PN-1000724), Xenium Cell Segmentation Staining Reagents (PN-1000661), and Xenium Prime 5K decoding reagents and consumables according to the manufacturer’s instructions, unless otherwise noted. No custom add-on panel was used.

#### Xenium sample preparation

FFPE blocks were sectioned at 5 µm and mounted within the sample area of Xenium slides. Slides were equilibrated to room temperature for 30 min, air-dried overnight at room temperature, baked at 42 °C for 3 h, and stored in a desiccated environment until processing. Immediately before deparaffinization, slides were baked at 60 °C for 120 min. Slides were then deparaffinized by sequential immersion in xylene (2 × 10 min), 100% ethanol (2 × 3 min), 95% ethanol (2 × 3 min), 70% ethanol (1 × 3 min), and nuclease-free water (20 s). After rehydration, slides were immediately assembled into Xenium Prime cassettes to prevent tissue drying. Tissue sections were decrosslinked using Xenium Prime Sample Prep Reagents according to the manufacturer’s protocol, including incubation at 80 °C for 30 min. Sections then underwent priming hybridization at 50 °C for 90 min, followed by a post-priming wash at 50 °C for 30 min, RNase treatment at 37 °C for 20 min, and polishing at 37 °C for 1 h. Probe hybridization with the Xenium Prime 5K Human Pan Tissue and Pathways Panel was performed at 50 °C for 16 h.

On the following day, sections were subjected to a post-hybridization wash at 35 °C for 15 min, ligation at 42 °C for 30 min, amplification enhancement at 4 °C for 2 h, and amplification at 30 °C for 90 min. For cell segmentation, amplified sections were processed using Xenium Cell Segmentation Staining Reagents. Sections were equilibrated through 70% ethanol, 100% ethanol, 100% ethanol, 70% ethanol, and PBS-T, blocked in diluted Xenium Block and Stain Buffer for 1 h at room temperature, and incubated with Xenium Multi-Tissue Stain Mix overnight at 4 °C. On the following day, sections underwent stain enhancement for 20 min at room temperature, autofluorescence quenching with diluted Reducing Agent B for 10 min followed by AF Solution for 10 min, drying at 37 °C for 5 min, and nuclei staining for 1 min at room temperature. Slides were then loaded onto the Xenium Analyzer with the appropriate decoding reagents and consumables, and imaging, transcript decoding, and cell segmentation were performed according to the manufacturer’s instructions.

#### Post-Xenium H&E staining

After Xenium imaging, slides were removed from the Xenium cassette and processed for hematoxylin and eosin (H&E) staining. Residual quencher was removed by incubating slides for 10 min at room temperature in freshly prepared quencher removal solution consisting of 69.6 mg sodium hydrosulfite dissolved in 40 mL molecular-grade water, followed by three 1-min washes in molecular-grade water. Slides were either processed immediately for H&E staining or stored in 1× PBS at 4 °C for up to 2 days before staining. For H&E staining, slides were immersed in molecular-grade water for 2 min, stained in Mayer’s hematoxylin for 20 min, washed in molecular-grade water three times for 1 min each, differentiated in Dako Bluing Solution for 1 min, and rinsed in molecular-grade water for 1 min. Sections were then dehydrated through 70% ethanol for 3 min and 95% ethanol for 3 min, stained in Eosin Y, alcoholic for 4.5 min, and further dehydrated through 95% ethanol (2 × 30 s) and 100% ethanol (2 × 30 s). Slides were cleared in xylene (2 × 3 min), coverslipped, and dried in a fume hood for 30 min. H&E-stained slides were scanned at 20× magnification using a Leica Aperio CS2 whole-slide scanner.

### Co-culture cytotoxicity experiment

A cytotoxicity experiment was conducted where GFP-tagged Py230, 4T1, and MDA-MB-231 cancer cells were co-cultured with NK cells isolated from the spleen of C57BL/6J mice at a 1:1 and 2:1 ratio in the presence and absence of S100A4-enriched CAF CM. The plate was incubated for 24 h in a 5% CO2 atmosphere at 37°C. After 24 h in culture, images were captured by an Echo Rebel microscope (Echo, San Diego, CA, USA), and the cell count was analyzed for cytotoxicity effects.

### Co-culture spheroid experiment

This experiment was performed using Nunclon™ Sphera™ 96-Well, Nunclon Sphera-Treated, U-Shaped-Bottom Microplates (Thermo Fisher Scientific; Waltham, MA, USA) to assess the cytotoxicity effect of NK cells on GFP-tagged Py230 spheroids in the presence and absence of S100A4-enriched CAF CM. A single-cell suspension was combined with Matrigel matrix gel at a 1:1 ratio, and the plate was incubated for 5-7 days in a 5% CO2 atmosphere at 37°C. After 5-7 days in culture, spheroid images were captured by an Echo Rebel microscope (Echo, San Diego, CA, USA).

### Proteome Profiler Mouse Cytokine Array

Filtered supernatant of IL-12 p70 Recombinant Protein (PeproTech, #210-12-10UG, Cranbury, NJ, USA) and Mouse IL-15 Recombinant Protein (PeproTech, #210-15-10UG, Cranbury, NJ, USA)-activated spleen NK cells incubated in the presence and absence of S100A4-enriched CAF CM for 24h and these samples were utilized by a Proteome Profiler Mouse Cytokine Array Kit, Panel A (R&D Systems, # ARY006, Minneapolis, MN, USA). Images of the blots were taken with a ChemiDoc™ MP Imaging System (Bio-Rad, # 12003154, Hercules, CA, USA).

### 3D Co-culture and CM experiment

This experiment was performed using an 8-well Nunc™ Lab-Tek™ Chamber Slide System (Thermo Fisher Scientific, #177445; Waltham, MA, USA) coated with Matrigel matrix gel to assess the cytotoxic effect of NK cells on GFP-tagged Py230 cells in the presence and absence of S100A4-enriched CAF CM and co-cultured S100A4-enriched CAF cells. A single-cell suspension was combined with Matrigel matrix gel at a 1:1 ratio, and the plate was incubated for 5-7 days in a 5% CO2 atmosphere at 37°C. After 7 days in culture, images of the 3D co-culture were captured by an Echo Rebel microscope (Echo, San Diego, CA, USA).

### CAF inhibition *in vitro* experiment

Niclosamide^69^ (25g, Cayman Chemical, #10649, Ann Arbor, MI, USA) was utilized at a concentration of 1.5 μM to inhibit S100a4-enriched cells. UAMC1110^70^ (10 mM, MedChemExpress, #HY-100684, Monmouth Junction, NJ, USA) was utilized at a concentration of 1 μM to inhibit Fibroblast Activation Protein (FAP)-enriched cells. gp38 InVivoMAb^71^ anti-mouse Podoplanin (gp38) Clone 8.1.1 (1mg, Bio X Cell, #BE0236, Lebanon, NH, USA) was utilized at a concentration of 1 μM to inhibit Podoplanin (PDPN)-enriched cells. The inhibition effects were analyzed by Western blot. *β-actin* was used as a housekeeping gene.

### BCAT1 *in vitro* inhibition experiment

Cells were pretreated with ERG240^70^ (60 μM, TargetMol, #1415683-79-2, Wellesley Hills, MA, USA), an inhibitor of BCAT1, for 24 h, and anchorage-independent growth of cancer stem cells was assessed. Single-cell suspension was seeded in low-attachment plates and incubated for 5-7 days in a 5% CO2 atmosphere at 37°C. After 7 days in culture, images of the 3D co-culture were captured by Echo Rebel microscope (Echo, San Diego, CA, USA). The inhibition effects were analyzed by Western blot. β-actin was used as a housekeeping gene.

### Real-time PCR

Total RNA was extracted from cultured cells and frozen tumors using Quick-RNA™ MiniPrep (Zymo Research, # R1055, Irvine, CA, USA), and the purity was analyzed by DeNovix DS-11 Series nanodrop Spectrophotometer (DeNovix). RNA from each sample was reverse transcribed using the Applied Biosystems High-Capacity cDNA Reverse Transcription Kit (Bio-Rad Laboratories, #43-688-14, Hercules, CA, USA). Analysis for gene expression by quantitative real-time PCR was performed^29^ using SsoAdvanced™ Universal SYBR Green Supermix (Fisher Scientific, #1725271, Waltham, MA, USA). *Gapdh* was used as a housekeeping gene.

### Western Blot

Total protein samples were extracted from cultured cells and frozen tumors using the RIPA Lysis Buffer System (Santa Cruz Biotechnology Inc., # sc-24948A, Dallas, TX, USA). Samples were lysed on ice for 30min and then centrifuged for 20min at 12,000 rpm at 4°C. Equal amounts of protein were resolved by 12% or 15% SDS–PAGE. Transferred proteins were stained with primary antibodies, followed by an HRP-conjugated secondary antibody. β-actin was used as a housekeeping gene^65^.

### ^13^C glucose labeling experiment

TRACER experiment, in collaboration with MD Anderson Cancer Center^43^, in which S100A4-enriched CAF cells were incubated in media (10% FBS (dialyzed, Thermo Scientific), 2 mM L-glutamine (GIBCO), 2.5 mM 13C6 glucose in glucose-free DMEM) containing uniformly 13C-labeled glucose ([U-13C] glucose for 24-48 h to allow labeling of all glucose-derived metabolites in the cells. After 24-48 h, the cells were washed with PBS to remove any residual [U-13C] glucose before incubation with media containing unlabeled glucose (10% FBS, 2 mM L-glutamine, 2.5 mM glucose in DMEM) to collect the CM, which contained labeled secreted metabolites. Additionally, the CM was fed to various cell types (Py230 cancer cells and NK cells), and after incubation in media containing [U-13C] glucose, the cells were washed with PBS to remove any residual [U-13C] glucose before incubation with media containing unlabeled glucose (10% FBS, 2 mM L-glutamine, 2.5 mM glucose in DMEM) to collect the cells, which contained labeled secreted metabolites. The CM and fed cell samples’ metabolites were extracted according to a standard protocol (using ammonium bicarbonate and ammonium hydroxide in 80% MEOH, respectively) and measured for metabolomic profiling.

### ELISA Assay

ELISA MAX™ Deluxe Set Mouse IFN-γ (BioLegend, #430804, San Diego, CA, USA) was utilized to measure IFN-γ levels in cell culture supernatants^65^. The cell culture supernatant of Mouse IL-12 p70 Recombinant Protein (PeproTech, #210-12-10UG, Cranbury, NJ, USA) and Mouse IL-15 Recombinant Protein (PeproTech, #210-15-10UG, Cranbury, NJ, USA)-activated spleen NK cells under various incubation conditions, utilizing BCAAs: valine, leucine, and isoleucine (gifted by Guoyao Wu, TAMU Department of Animal Science), and BCKAs: 3-Methyl-2-oxobutanoic acid (keto-valine, MedChemExpress, #HY-W006057, Monmouth Junction, NJ), 3-Methyl-2-oxopentanoic acid sodium salt (keto-isoleucine, MilliporeSigma, #198978-5G, Burlington, MA, USA), and α-Ketoisocaproic Acid (keto-leucine, Cayman Chemical, # 34749, Ann Arbor, MI). The absorbance was recorded at 450nm using a spectrophotometric plate reader (Molecular Devices; San Jose, CA, USA) to analyze IFN-γ cytokine. ELISA MAX™ Deluxe Set Mouse IFN-γ (BioLegend, #430804, San Diego, CA, USA) was utilized to measure IFN-γ levels in cell culture supernatants. The cell culture supernatant of non-activated and Mouse IL-12 p70 Recombinant Protein (PeproTech, #210-12-10UG, Cranbury, NJ, USA) and Mouse IL-15 Recombinant Protein (PeproTech, #210-15-10UG, Cranbury, NJ, USA)-activated spleen NK cells under various incubation conditions, utilizing CAF CM, BCAT1 inhibitor, ERG240 (60 μM, TargetMol, #1415683- 79-2, Wellesley Hills, MA, USA), and BCAT2 inhibitor, BCATc Inhibitor 2 (Cayman Chemical, #9002002, Ann Arbor, MI). The absorbance was recorded at 450nm using a spectrophotometric plate reader (Molecular Devices; San Jose, CA, USA) to analyze IFN-γ cytokine.

### Targeted (amino acid) metabolomics

Sample Extraction: NF CM and CAF CM (n=3) were weighed and extracted with a methanol-based extraction method. Briefly, 800 uL ice ice-cold methanol was added to samples in a bead-based lysis tube (Bertin, Rockville, MD). Samples were extracted on a Precyllys 24 (Bertin) tissue homogenizer for 30 seconds at a speed of 6000. The supernatant was collected and passed through a 0.2 um nylon filter (Merck Millipore, Burlington, MA). 500 uL of the filtered aqueous phase was then passed through a 3 kDa cutoff column (Thermo Scientific), and the flow-through was collected for analysis.

Sample Analysis: Untargeted liquid chromatography high-resolution accurate mass spectrometry (LC-HRAM) analysis was performed on a Q Exactive Plus Orbitrap mass spectrometer (Thermo Scientific, Waltham, MA) coupled to a binary pump HPLC (UltiMate 3000, Thermo Scientific). Full MS spectra were obtained at 70,000 resolution (200 m/z) with a scan range of 50-750 m/z. Full MS followed by ddMS2 scans were obtained at 35,000 resolution (MS1) and 17,500 resolution (MS2) with a 1.5 m/z isolation window and a stepped NCE (20, 40, 60). Samples were maintained at 4 °C before injection. The injection volume was 10 µL. Chromatographic separation was achieved on a Synergi Fusion 4µm, 150 mm x 2 mm reverse-phase column (Phenomenex, Torrance, CA) maintained at 30 °C using a solvent gradient method. Solvent A was water (0.1% formic acid). Solvent B was methanol (0.1% formic acid). The gradient method used was 0-5 min (10% B to 40% B), 5-7 min (40% B to 95% B), 7-9 min (95% B), 9-9.1 min (95% B to 10% B), 9.1-13 min (10% B). The flow rate was 0.4 mL min^-1^. Sample acquisition was performed Xcalibur (Thermo Scientific). Data analysis was performed with Compound Discoverer 3.3 (Thermo Scientific).

### Branched-Chain Amino Acid (BCAA) Analysis

Amino acid concentrations were determined by pre-column derivatization with o-phthalaldehyde (OPA) followed by reverse-phase HPLC with fluorescence detection. Samples were deproteinized by adding 40 µL of 1.5 M perchloric acid to 40 µL of sample on ice. After vortexing for 15 s, 20 µL of ice-cold 2 M potassium carbonate was added for neutralization, and samples were vortexed for an additional 15 s. Samples were centrifuged at 10,000 rpm for 5 min, and the resulting supernatants were collected for analysis. HPLC-grade water and amino acid standards (Agilent Technologies, 5061-3331) were processed using the same procedure as the blank and calibration standards, respectively. For amino acid quantification, 20 µL of sample supernatant was mixed with 20 µL of internal standard and analyzed using an Agilent 1260 Infinity Series HPLC system controlled by OpenLab CDS software. Amino acids were derivatized using an OPA reagent (Agilent Technologies, 5061-3335) and separated using reverse-phase chromatography on a Thermo Scientific ODS Hypersil C18 column. Detection was performed using an Agilent fluorescence detector (G1321B) with excitation and emission wavelengths of 340 nm and 450 nm, respectively. Chromatographic separation was performed at a flow rate of 0.45 mL/min using a gradient elution program beginning with 100% mobile phase A at 0 min and reaching 60% mobile phase B at 17 min.

### Branched-Chain Keto Acid (BCKA) Analysis

Branched-chain keto acids (BCKAs) were quantified using o-phenylenediamine (OPD) derivatization followed by HPLC analysis. Samples were deproteinized by adding 40 µL of 1.5 M perchloric acid to 40 µL of sample on ice, followed by vortex mixing for 15 s and centrifugation at 10,000 rpm for 5 min. A 50 µL aliquot of supernatant was reacted with 50 µL of 25 mM o-phenylenediamine prepared in 2 M HCl. Samples were incubated at 100°C for 30 min while protected from light and subsequently cooled on ice for 5 min. The resulting OPD-derived products were analyzed by HPLC without additional derivatization. Chromatographic separation was performed using the following mobile phases: mobile phase A consisted of 20 mM sodium acetate buffer containing 0.018% (v/v) triethylamine, 0.05 mM EDTA, and 0.3% (v/v) tetrahydrofuran (THF), adjusted to pH 7.2. Mobile phase B consisted of 20% 100 mM sodium acetate buffer, 40% acetonitrile, and 40% methanol (v/v).

### Orthotopic injection of breast cancer cells

Orthotopic injection of breast cancer cells into the mammary fat pad was performed. On the day of operation, the Py230 cells derived from C57BL/6J mice were washed once with phosphate-buffered saline (PBS) and trypsinized. Trypsin was quenched by adding 4 mL of F12K complete medium. The cells were centrifuged at 300×g for 5 min to remove the serum, then resuspended in medium. Subsequently, the cells were counted to 1×10^6^ cells/mouse. The cells were resuspended in Matrigel and kept on ice. Female mice were anesthetized using 3% inhalant isoflurane. The mammary fat pad was squeezed using tweezers to expose the fat pad for the injection. Py230 cells were then injected using an insulin syringe with a volume of 100 μL. The length and width of the tumor mass were measured every 2-3 days using slide calipers, and tumor volume was calculated as follows: tumor volume = length of the tumor x (width of the tumor) 2 / 2 (mm^3^), and tumor burden = tumor weight/mouse weight x 100.

### Co-injection: CAF & NF cells

Py230 cancer cells (500,000) were co-injected at a 1:1 ratio with 500,000 S100A4-enriched CAF cells or with NF cells. The cells were resuspended in Matrigel and kept on ice. Female mice were anesthetized using 3% inhalant isoflurane. The mammary fat pad was squeezed using tweezers to expose the fat pad for the injection. Py230 cells were then injected using an insulin syringe with a volume of 100 μL. The length and width of the tumor mass were measured every 2-3 days using slide calipers, and tumor volume and tumor burden were measured to analyze the co-injection effects.

### Niclosamide inhibition *in vivo* experiment

Py230 cells (1×10^6^ cells) were injected using an insulin syringe with a volume of 100 μL. Once palpable tumors developed, Niclosamide^69^ (25g, Cayman Chemical, #10649, Ann Arbor, MI, USA), an inhibitor of S100A4, was administered at 20 mg/kg/day (DMSO: Tween-80:H2O= 3:4:8) to C57BL/6J mice injected by intraperitoneal injection for 4 weeks. The length and width of the tumor mass were measured every 2-3 days using slide calipers, and tumor volume and tumor burden were measured to analyze the inhibition effects.

### ERG240 *in vivo* inhibition experiment

Py230 cells (1×10^6^ cells) were injected using an insulin syringe with a volume of 100 μL. Once palpable tumors developed, ERG240 (5 mg/kg/day, TargetMol, #1415683-79-2, Wellesley Hills, MA, USA), an inhibitor of BCAT1^72^, was administered to C57BL/6J mice injected by intraperitoneal injection for 4 weeks. The length and width of the tumor mass were measured every 2-3 days using slide calipers, and tumor volume and tumor burden were measured to analyze the inhibition effects.

### NK depletion *in vivo* experiment

Py230 cells (1×10^6^ cells) were injected using an insulin syringe with a volume of 100 μL. Once palpable tumors developed, MAb anti-mouse NK1.1, clone PK136^73^ (200μg, Bio X Cell, #BE0036, Lebanon, NH, USA), was administered to C57BL/6J mice for *in vivo* depletion of NK1.1-expressing cells. InVivoMAb mouse IgG2a isotype control (200μg, Bio X Cell, # BE0085, Lebanon, NH, USA) was administered to C57BL/6J mice. Mice were injected by intraperitoneal injection every third day. Additionally, InVivoPure pH 7.0 Dilution Buffer (Bio X Cell, #IP0070, Lebanon, NH, USA) was utilized. The length and width of the tumor mass were measured every 2-3 days using slide calipers, and tumor volume and tumor burden were measured to analyze the depletion effects.

### Statistical Analysis

Data collected were analyzed using FlowJo software (BD Biosciences) and GraphPad Prism. UMAPs generated in FlowJo. Pairs of samples were compared using an unpaired t-test. When comparing more than two sets of data, statistical significance was determined by either one-way ANOVA with Tukey’s multiple comparison test or two-way ANOVA with Sidak’s multiple comparisons test. All reported values represent the mean and standard error of the mean from three to nine biological replicates, with at least technical triplicates within each experiment. Whenever possible, all data points are included in graphical representations of data. All graphs show mean ± SEM unless stated otherwise. Statistical significance was accepted when p<0.05, and “ns” in all figures indicates p-values greater than 0.05. All details were shown in the figure legends.

## References

1 Hanahan, D. & Coussens, L. M. Accessories to the crime: functions of cells recruited to the tumor microenvironment. Cancer Cell 21, 309–322 (2012). 10.1016/j.ccr.2012.02.022

2 Balkwill FR, C. M., Hagemann T The tumor microenvironment at a glance. J Cell Sci. J Cell Sci 125, 5591–5596 (2012).

3 Quail, D. F. & Joyce, J. A. Microenvironmental regulation of tumor progression and metastasis. Nat Med 19, 1423–1437 (2013). 10.1038/nm.3394

4 Denkert, C. et al. Clinical and molecular characteristics of HER2-low-positive breast cancer: pooled analysis of individual patient data from four prospective, neoadjuvant clinical trials. Lancet Oncol 22, 1151–1161 (2021). 10.1016/S1470-2045(21)00301-6

5 Modi, S. et al. Trastuzumab Deruxtecan in Previously Treated HER2-Low Advanced Breast Cancer. N Engl J Med 387, 9–20 (2022). 10.1056/NEJMoa2203690

6 Schettini, F. et al. Author Correction: Clinical, pathological, and PAM50 gene expression features of HER2-low breast cancer. NPJ Breast Cancer 9, 32 (2023). 10.1038/s41523-023-00538-x

7 Tarantino, P. et al. HER2-Low Breast Cancer: Pathological and Clinical Landscape. J Clin Oncol 38, 1951–1962 (2020). 10.1200/JCO.19.02488

8 Kalluri, R. The biology and function of fibroblasts in cancer. Nat Rev Cancer 16, 582–598 (2016). 10.1038/nrc.2016.73

9 Bartoschek, M. et al. Spatially and functionally distinct subclasses of breast cancer-associated fibroblasts revealed by single cell RNA sequencing. Nat Commun 9, 5150 (2018). 10.1038/s41467-018-07582-3

10 Sahai E, e. a. A framework for advancing our understanding of cancer-associated fibroblasts. Nat Rev Cancer Nature reviews Cancer 20, 174-186 (2020).

11 Chen, Y., McAndrews, K. M. & Kalluri, R. Clinical and therapeutic relevance of cancer-associated fibroblasts. Nat Rev Clin Oncol 18, 792–804 (2021). 10.1038/s41571-021-00546-5

12 Orimo, A. et al. Stromal fibroblasts present in invasive human breast carcinomas promote tumor growth and angiogenesis through elevated SDF-1/CXCL12 secretion. Cell 121, 335–348 (2005). 10.1016/j.cell.2005.02.034

13 Ohlund, D., Elyada, E. & Tuveson, D. Fibroblast heterogeneity in the cancer wound. J Exp Med 211, 1503–1523 (2014). 10.1084/jem.20140692

14 Fiori, M. E. et al. Cancer-associated fibroblasts as abettors of tumor progression at the crossroads of EMT and therapy resistance. Mol Cancer 18, 70 (2019). 10.1186/s12943-019-0994-2

15 Bu, L., Baba, H., Yasuda, T., Uchihara, T. & Ishimoto, T. Functional diversity of cancer-associated fibroblasts in modulating drug resistance. Cancer Sci 111, 3468–3477 (2020). 10.1111/cas.14578

16 Costa, A. et al. Fibroblast Heterogeneity and Immunosuppressive Environment in Human Breast Cancer. Cancer Cell 33, 463–479 e410 (2018). 10.1016/j.ccell.2018.01.011

17 Kieffer, Y. et al. Single-Cell Analysis Reveals Fibroblast Clusters Linked to Immunotherapy Resistance in Cancer. Cancer Discov 10, 1330–1351 (2020). 10.1158/2159-8290.CD-19-1384

18 Lambrechts, D. et al. Phenotype molding of stromal cells in the lung tumor microenvironment. Nat Med 24, 1277–1289 (2018). 10.1038/s41591-018-0096-5

19 Guillerey, C., Huntington, N. D. & Smyth, M. J. Targeting natural killer cells in cancer immunotherapy. Nat Immunol 17, 1025–1036 (2016). 10.1038/ni.3518

20 Vivier, E. et al. Innate or adaptive immunity? The example of natural killer cells. Science 331, 44–49 (2011). 10.1126/science.1198687

21 Mamessier, E. et al. Peripheral blood NK cells from breast cancer patients are tumor-induced composite subsets. J Immunol 190, 2424–2436 (2013). 10.4049/jimmunol.1200140

22 Krneta, T., Gillgrass, A., Chew, M. & Ashkar, A. A. The breast tumor microenvironment alters the phenotype and function of natural killer cells. Cell Mol Immunol 13, 628–639 (2016). 10.1038/cmi.2015.42

23 Mamessier, E. et al. Human breast cancer cells enhance self tolerance by promoting evasion from NK cell antitumor immunity. J Clin Invest 121, 3609–3622 (2011). 10.1172/JCI45816

24 Balsamo, M. et al. Melanoma-associated fibroblasts modulate NK cell phenotype and antitumor cytotoxicity. Proc Natl Acad Sci U S A 106, 20847–20852 (2009). 10.1073/pnas.0906481106

25 Savas, P., et al. Publisher Correction: Single-cell profiling of breast cancer T cells reveals a tissue-resident memory subset associated with improved prognosis. Nat Med 24, 1941 (2018). 10.1038/s41591-018-0176-6

26 Francescone, R. et al. Netrin G1 Promotes Pancreatic Tumorigenesis through Cancer-Associated Fibroblast-Driven Nutritional Support and Immunosuppression. Cancer Discov 11, 446–479 (2021). 10.1158/2159-8290.CD-20-0775

27 Wong, T., Kang, R. & Yun, K. The multi-faceted immune modulatory role of S100A4 in cancer and chronic inflammatory disease. Front Immunol 16, 1525567 (2025). 10.3389/fimmu.2025.1525567

28 Bogachek, M. et al. S100A4/FSP1: A Prognostic Marker and a Promising Target for Antitumor Therapy. Int J Mol Sci 26 (2025). 10.3390/ijms26199370

29 Friedman, G. et al. Cancer-associated fibroblast compositions change with breast cancer progression linking the ratio of S100A4(+) and PDPN(+) CAFs to clinical outcome. Nat Cancer 1, 692–708 (2020). 10.1038/s43018-020-0082-y

30 Croizer, H. et al. Deciphering the spatial landscape and plasticity of immunosuppressive fibroblasts in breast cancer. Nat Commun 15, 2806 (2024). 10.1038/s41467-024-47068-z

31 Ireland, L. et al. Blockade of insulin-like growth factors increases efficacy of paclitaxel in metastatic breast cancer. Oncogene 37, 2022–2036 (2018). 10.1038/s41388-017-0115-x

32 Guo, Q. et al. Physiologically activated mammary fibroblasts promote postpartum mammary cancer. JCI Insight 2, e89206 (2017). 10.1172/jci.insight.89206

33 Sharon, Y., Alon, L., Glanz, S., Servais, C. & Erez, N. Isolation of normal and cancer-associated fibroblasts from fresh tissues by Fluorescence Activated Cell Sorting (FACS). J Vis Exp, e4425 (2013). 10.3791/4425

34 Liu, C. et al. The conditioned medium from mesenchymal stromal cells pretreated with proinflammatory cytokines promote fibroblasts migration and activation. PLoS One 17, e0265049 (2022). 10.1371/journal.pone.0265049

35 Chen, C. Y. et al. Cancer-Associated-Fibroblast-Mediated Paracrine and Autocrine SDF-1/CXCR4 Signaling Promotes Stemness and Aggressiveness of Colorectal Cancers. Cells 13 (2024). 10.3390/cells13161334

36 Strating, E. et al. Co-cultures of colon cancer cells and cancer-associated fibroblasts recapitulate the aggressive features of mesenchymal-like colon cancer. Front Immunol 14, 1053920 (2023). 10.3389/fimmu.2023.1053920

37 Johansson, A. C. et al. Cancer-associated fibroblasts induce matrix metalloproteinase-mediated cetuximab resistance in head and neck squamous cell carcinoma cells. Mol Cancer Res 10, 1158–1168 (2012). 10.1158/1541-7786.MCR-12-0030

38 Milani, M. et al. Targeting S100A4 with niclosamide attenuates inflammatory and profibrotic pathways in models of amyotrophic lateral sclerosis. J Neuroinflammation 18, 132 (2021). 10.1186/s12974-021-02184-1

39 Yao, L. et al. Cancer-associated fibroblasts impair the cytotoxic function of NK cells in gastric cancer by inducing ferroptosis via iron regulation. Redox Biol 67, 102923 (2023). 10.1016/j.redox.2023.102923

40 Wu, S. Z. et al. A single-cell and spatially resolved atlas of human breast cancers. Nat Genet 53, 1334–1347 (2021). 10.1038/s41588-021-00911-1

41 Michel, T. et al. Mouse lung and spleen natural killer cells have phenotypic and functional differences, in part influenced by macrophages. PLoS One 7, e51230 (2012). 10.1371/journal.pone.0051230

42 Aquino-Lopez, A., Senyukov, V. V., Vlasic, Z., Kleinerman, E. S. & Lee, D. A. Interferon Gamma Induces Changes in Natural Killer (NK) Cell Ligand Expression and Alters NK Cell-Mediated Lysis of Pediatric Cancer Cell Lines. Front Immunol 8, 391 (2017). 10.3389/fimmu.2017.00391

43 Becker, L. M. et al. Epigenetic Reprogramming of Cancer-Associated Fibroblasts Deregulates Glucose Metabolism and Facilitates Progression of Breast Cancer. Cell Rep 31, 107701 (2020). 10.1016/j.celrep.2020.107701

44 Wang, X. et al. Colorectal cancer cells establish metabolic reprogramming with cancer-associated fibroblasts (CAFs) through lactate shuttle to enhance invasion, migration, and angiogenesis. Int Immunopharmacol 143, 113470 (2024). 10.1016/j.intimp.2024.113470

45 Dimou, A., Tsimihodimos, V. & Bairaktari, E. The Critical Role of the Branched Chain Amino Acids (BCAAs) Catabolism-Regulating Enzymes, Branched-Chain Aminotransferase (BCAT) and Branched-Chain alpha-Keto Acid Dehydrogenase (BCKD), in Human Pathophysiology. Int J Mol Sci 23 (2022). 10.3390/ijms23074022

46 Wolfe, R. R. Branched-chain amino acids and muscle protein synthesis in humans: myth or reality? J Int Soc Sports Nutr 14, 30 (2017). 10.1186/s12970-017-0184-9

47 Ananieva, E. A. & Wilkinson, A. C. Branched-chain amino acid metabolism in cancer. Curr Opin Clin Nutr Metab Care 21, 64–70 (2018). 10.1097/MCO.0000000000000430

48 Peng, H., Wang, Y. & Luo, W. Multifaceted role of branched-chain amino acid metabolism in cancer. Oncogene 39, 6747–6756 (2020). 10.1038/s41388-020-01480-z

49 Selwan, E. M. & Edinger, A. L. Branched chain amino acid metabolism and cancer: the importance of keeping things in context. Transl Cancer Res 6, S578–S584 (2017). 10.21037/tcr.2017.05.05

50 Zhang, W. et al. Branched-chain amino acid transaminases as promising targets in tumor therapy. Front Cell Dev Biol 14, 1712076 (2026). 10.3389/fcell.2026.1712076

51 Zhu, Z. et al. Tumour-reprogrammed stromal BCAT1 fuels branched-chain ketoacid dependency in stromal-rich PDAC tumours. Nat Metab 2, 775–792 (2020). 10.1038/s42255-020-0226-5

52 Domagala, J. et al. The Tumor Microenvironment-A Metabolic Obstacle to NK Cells’ Activity. Cancers (Basel) 12 (2020). 10.3390/cancers12123542

53 Ben-Shmuel, A. et al. Cancer-Associated Fibroblasts Serve as Decoys to Suppress NK Cell Anticancer Cytotoxicity in Breast Cancer. Cancer Discov 15, 1247–1269 (2025). 10.1158/2159-8290.CD-24-0131

54 Kang, Z. R. et al. Deficiency of BCAT2-mediated branched-chain amino acid catabolism promotes colorectal cancer development. Biochim Biophys Acta Mol Basis Dis 1870, 166941 (2024). 10.1016/j.bbadis.2023.166941

55 Papathanassiu, A. E. et al. BCAT1 controls metabolic reprogramming in activated human macrophages and is associated with inflammatory diseases. Nat Commun 8, 16040 (2017). 10.1038/ncomms16040

56 Sarkar, M. et al. Cancer-associated fibroblasts: The chief architect in the tumor microenvironment. Front Cell Dev Biol 11, 1089068 (2023). 10.3389/fcell.2023.1089068

57 Freeman, P. & Mielgo, A. Cancer-Associated Fibroblast Mediated Inhibition of CD8+ Cytotoxic T Cell Accumulation in Tumours: Mechanisms and Therapeutic Opportunities. Cancers (Basel) 12 (2020). 10.3390/cancers12092687

58 Chen, M. et al. CAFs and T cells interplay: The emergence of a new arena in cancer combat. Biomed Pharmacother 177, 117045 (2024). 10.1016/j.biopha.2024.117045

59 Hanley, C. J. & Thomas, G. J. T-cell tumour exclusion and immunotherapy resistance: a role for CAF targeting. Br J Cancer 123, 1353–1355 (2020). 10.1038/s41416-020-1020-6

60 Chen, S., Zhu, H. & Jounaidi, Y. Comprehensive snapshots of natural killer cells functions, signaling, molecular mechanisms and clinical utilization. Signal Transduct Target Ther 9, 302 (2024). 10.1038/s41392-024-02005-w

61 Russo, E. et al. NK Cell Anti-Tumor Surveillance in a Myeloid Cell-Shaped Environment. Front Immunol 12, 787116 (2021). 10.3389/fimmu.2021.787116

62 Luo, J., Xiang, X., Gong, G. & Jiang, L. Cancer-associated fibroblast-mediated immune evasion: molecular mechanisms of stromal-immune crosstalk in the tumor microenvironment. Front Immunol 16, 1617662 (2025). 10.3389/fimmu.2025.1617662

63 Jung, M. K., Okekunle, A. P., Lee, J. E., Sung, M. K. & Lim, Y. J. Role of Branched-chain Amino Acid Metabolism in Tumor Development and Progression. J Cancer Prev 26, 237–243 (2021). 10.15430/JCP.2021.26.4.237

64 Xu, E., Ji, B., Jin, K. & Chen, Y. Branched-chain amino acids catabolism and cancer progression: focus on therapeutic interventions. Front Oncol 13, 1220638 (2023). 10.3389/fonc.2023.1220638

65 Ogunlusi, O. et al. LILRB4 regulates circadian disruption-induced mammary tumorigenesis via non-canonical WNT signaling pathway. Oncogene 44, 4491–4504 (2025). 10.1038/s41388-025-03597-5

66 Finak, G. et al. MAST: a flexible statistical framework for assessing transcriptional changes and characterizing heterogeneity in single-cell RNA sequencing data. Genome Biol 16, 278 (2015). 10.1186/s13059-015-0844-5

67 Nguyen, T., Maniyar, A., Sarkar, M., Sarkar, T. R. & Neelgund, G. M. The Cytotoxicity of Carbon Nanotubes and Hydroxyapatite, and Graphene and Hydroxyapatite Nanocomposites against Breast Cancer Cells. Nanomaterials (Basel) 13 (2023). 10.3390/nano13030556

68 Manuel Iglesias J, B. I., Garcia-Garcia F, Leis O, Vazquez-Martin A, et al., Mammosphere Formation in Breast Carcinoma Cell Lines Depends upon Expression of E-cadherin. Plos One 8 (2013).

69 Yin, L. et al. Niclosamide sensitizes triple-negative breast cancer cells to ionizing radiation in association with the inhibition of Wnt/beta-catenin signaling. Oncotarget 7, 42126–42138 (2016). 10.18632/oncotarget.9704

70 Zhu, M. et al. Fibroblast Activation Protein Promotes Pulmonary Artery Hypertension via Activation of the PTEN/PI3K/Akt Pathway. J Cardiovasc Pharmacol 86, 391–407 (2025). 10.1097/FJC.0000000000001735

71 Astarita, J. L. et al. The CLEC-2-podoplanin axis controls the contractility of fibroblastic reticular cells and lymph node microarchitecture. Nat Immunol 16, 75–84 (2015). 10.1038/ni.3035

72 Chen, L. et al. NR4A1 deficiency promotes carotid plaque vulnerability by activating integrated stress response via targeting Bcat1. Cell Mol Life Sci 82, 91 (2025). 10.1007/s00018-025-05602-2

73 Dean, I. et al. Rapid functional impairment of natural killer cells following tumor entry limits anti-tumor immunity. Nat Commun 15, 683 (2024). 10.1038/s41467-024-44789-z

