## Supplementary material for "A stromal metabolic program suppresses NK-cell immunity to drive tumor progression in HER2-low breast cancer": Carter Supplementary doc

Table 1.

Clinical Breast Cancer Patient Data

| Patient ID | HER2 | ER/PR | Clinical Stage | Neoadjuvant |
|---|---|---|---|---|
| 6 | 1+ | <1 | IB | Dose Dense-Doxorubicin and Cyclophosphamide-Paclitaxel |
| 8 | 1+ | <1 | IIIB | Sacituzumab+Pembrolizumab |
| 10 | 1+ | 0 | IIB | KN-522; Doxorubicin and Cyclophosphamide |
| 11 | 2+/ISH+ | <1 | IIIB | Taxotere, Carboplatin, Herceptin, and Perjeta |
| 12 | 2+/ISH+ | <1 | IIB | Taxotere, Carboplatin, Herceptin, and Perjeta |
| 14 | 1+ | <1 | IIIB | KN-522; Doxorubicin and Cyclophosphamide |
| 18 | 2+/ISH- | <10 | IIIC | Doxorubicin and Cyclophosphamide Paclitaxel |
| 24 | 1+ | <10 | IIB | Carboplatin & Gemcitabine |
| 26 | 1+ | 0 | IIIC | Pembrolizumab combined with Carboplatin and Paclitaxel |

Supplementary Figure 1.

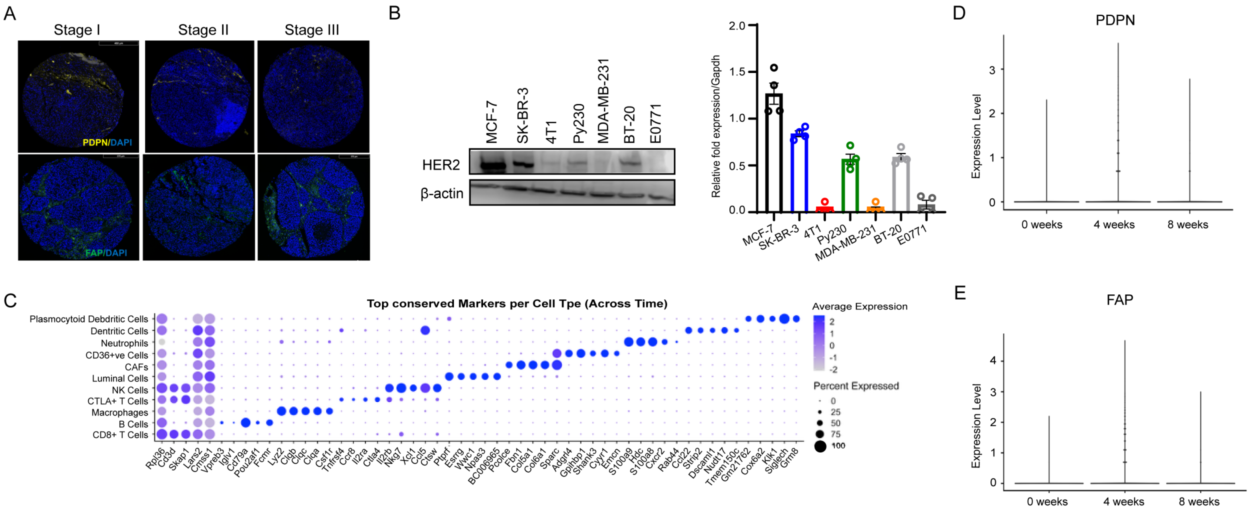

### Supplementary Figure 2.

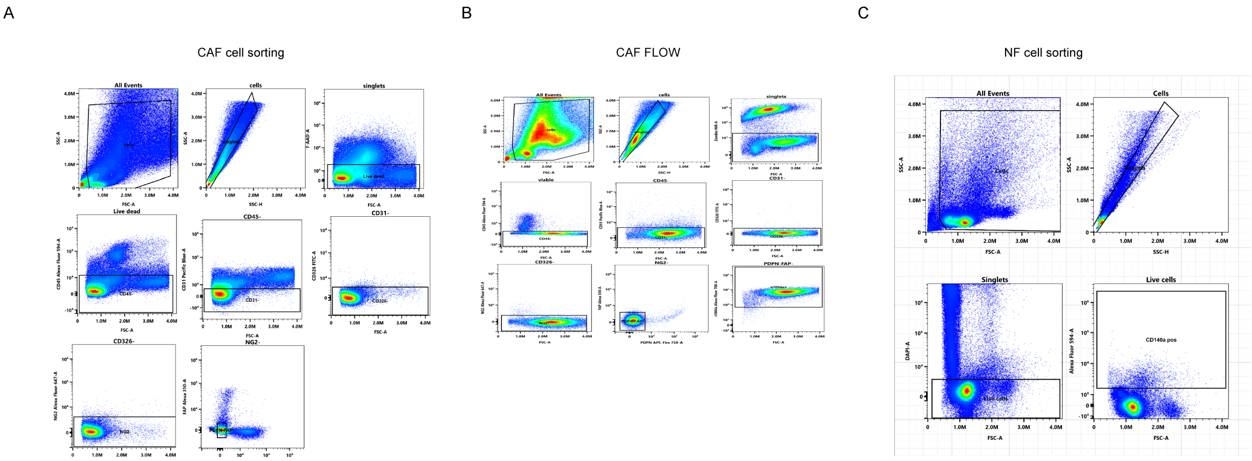


### Supplementary Figure 3.

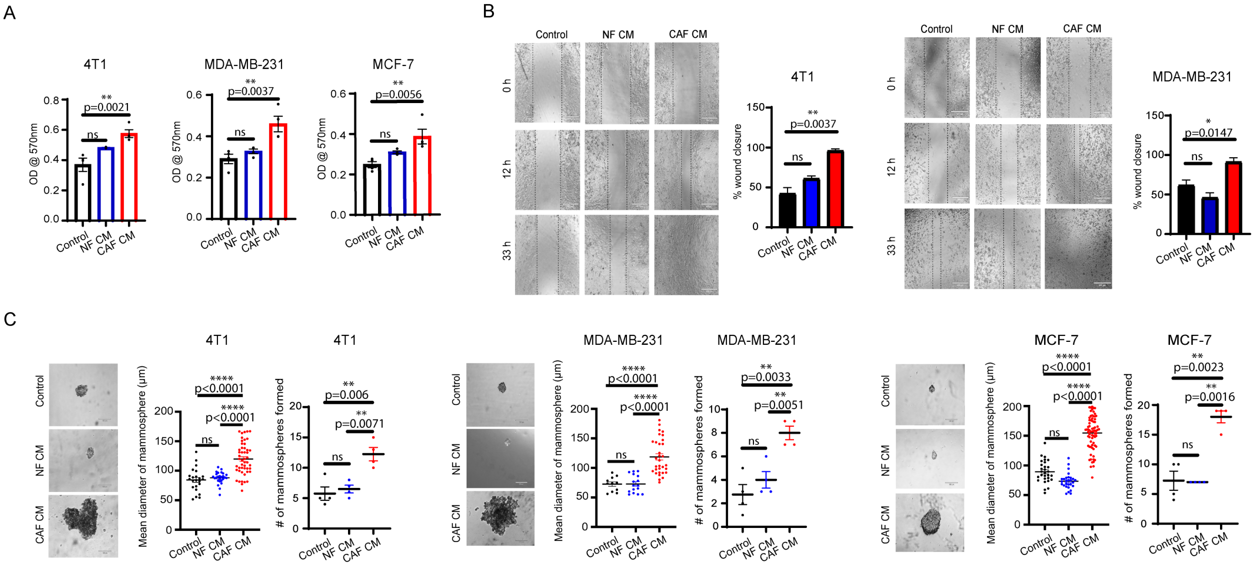

Supplementary Figure 4.

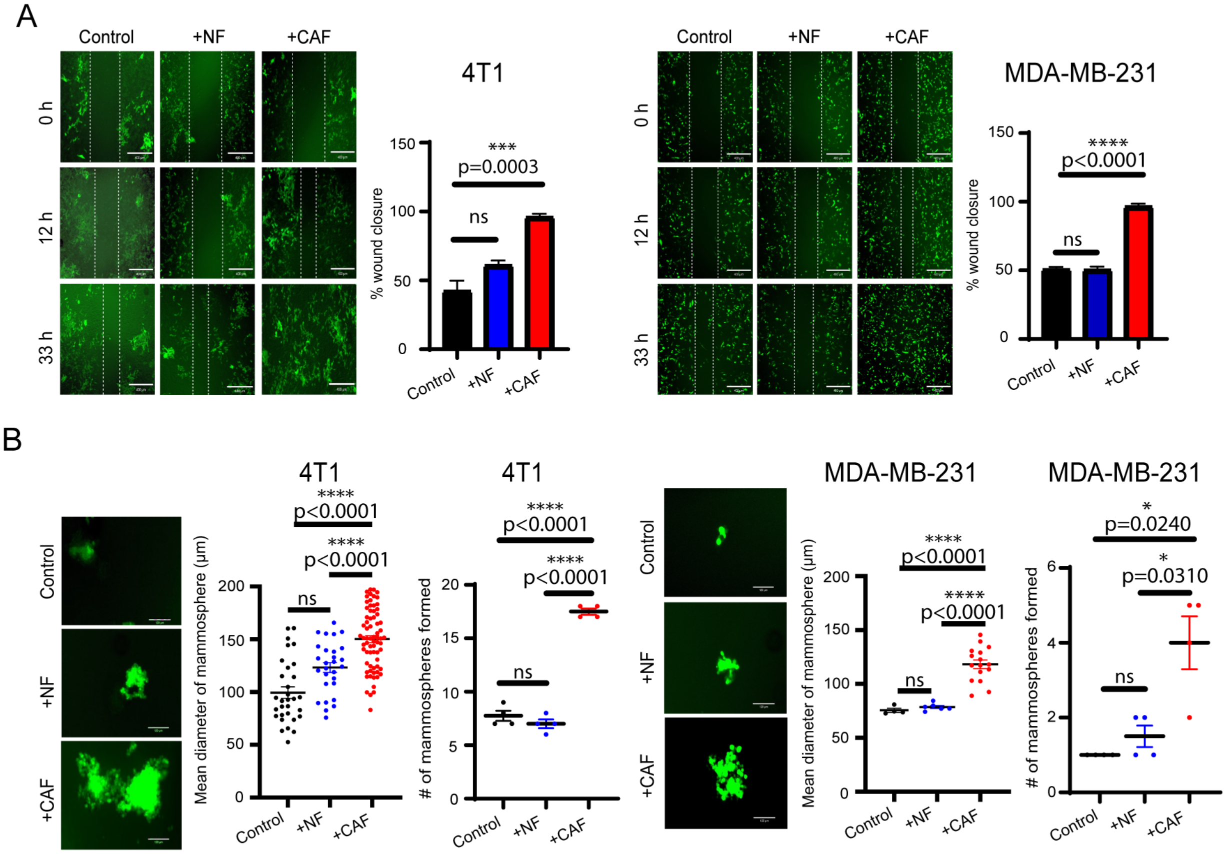

Supplementary Figure 5.

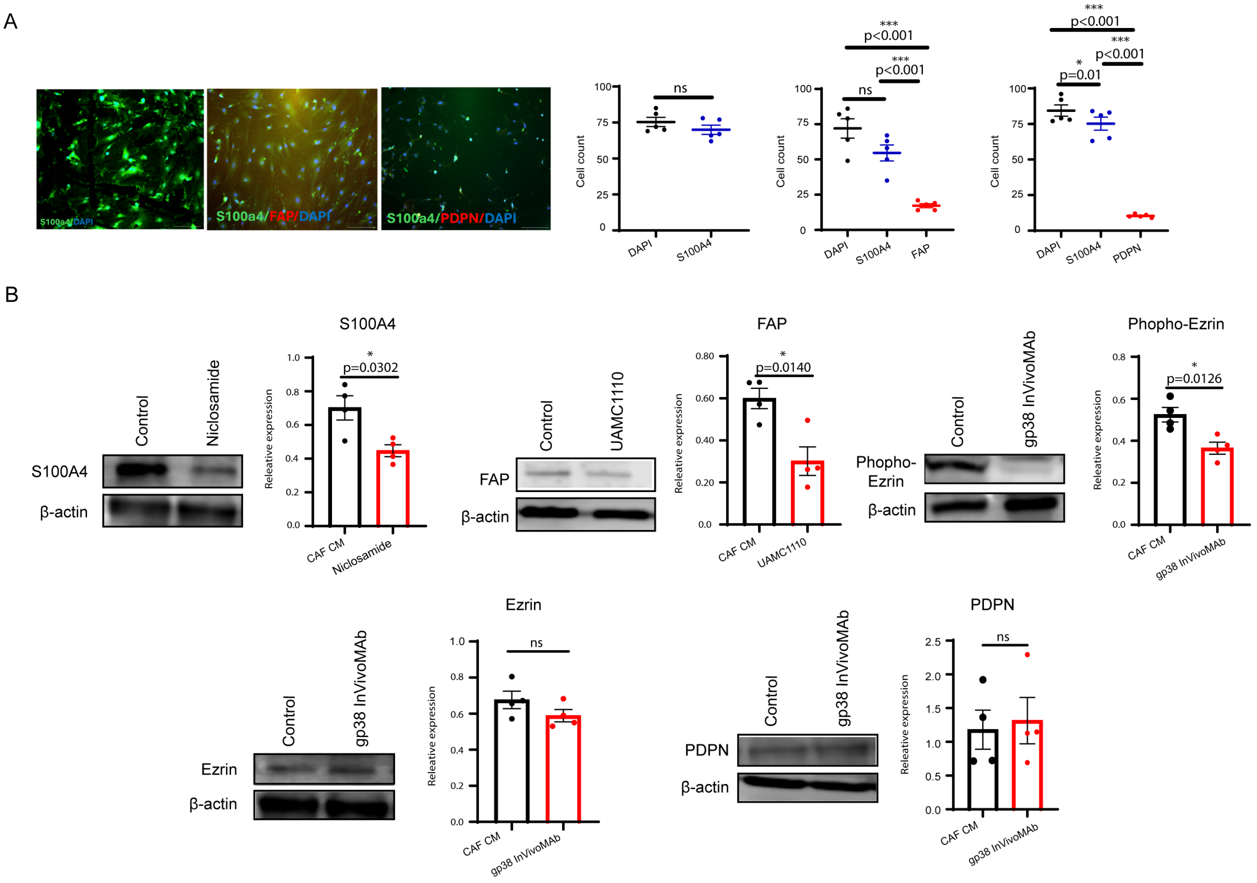

### Supplementary Figure 6.

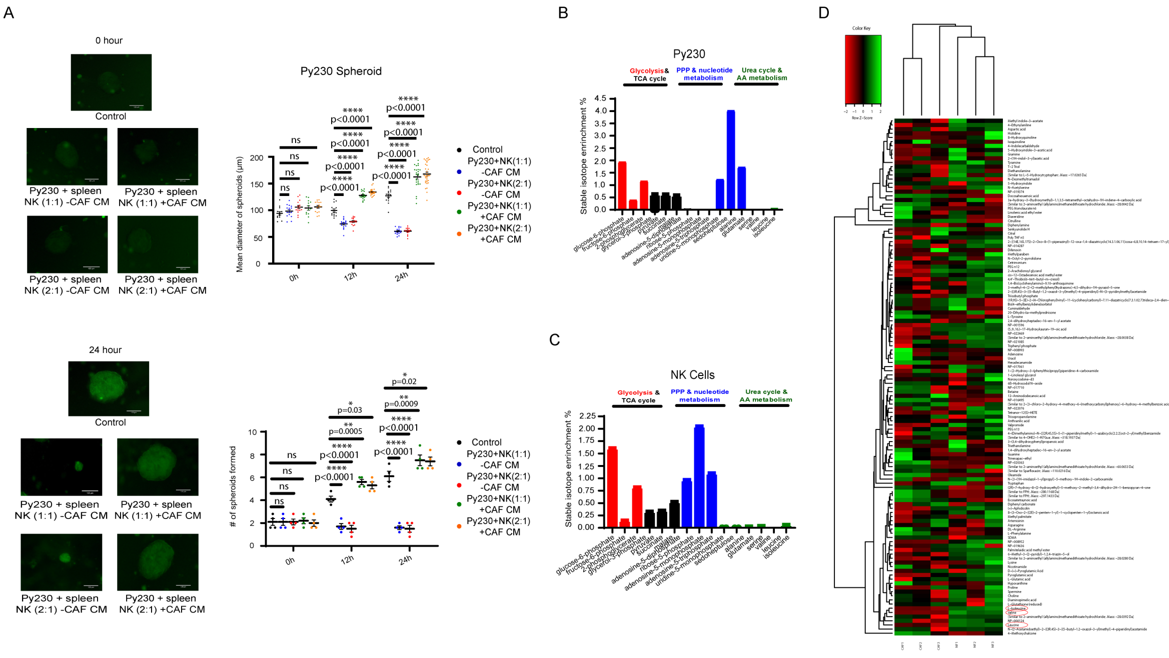


### Supplementary Figure 7.

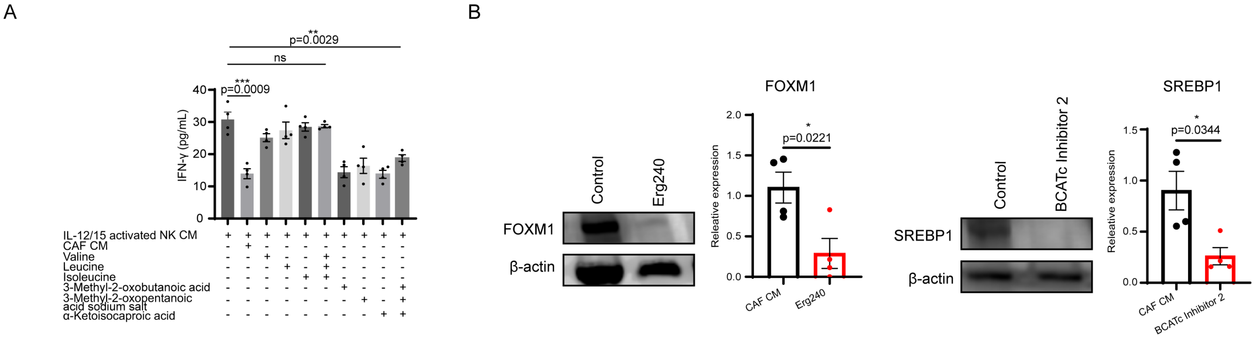

**Figure Legend:**

**Table 1: Clinical breast cancer patient data.**

Clinical breast cancer patient data including the following: patient ID, HER2 score, ER/PR, clinical stage, and neoadjuvant treatment.

**Supplementary Figure 1: PDPN and FAP CAFs enrichment during tumor progression.**

**(A)** Multiplex immunostaining of $PDPN^+$ and $FAP^+$ stromal cells from stage IA to stage IIIA. Scale bar: 450μm (top), 370μm (bottom). **(B)** Western blot analysis showing the expression of HER2 present in MCF-7, SK-BR-3, 4T1, Py230, MDA-MB-231, BT-20, and E0771 cancer cell lines. The densitometric analyses compare protein expression relative to β-actin (n = 4). **(C)** Top conserved markers per cell type via scRNA-seq. ScRNA-seq analysis showed an enrichment of **(D)** PDPN and **(E)** FAP CAFs during tumor progression.

**Supplementary Figure 2: S100A4+ cell collection via FACS.**

**(A)** Flow-activated cell sorting of the CAF population, gated on $CD45^-CD31^-CD326^-NG2^-PDPN^-FAP^-S100A4^+$. **(B)** Flow cytometry of the CAF population, gated on $CD45^-CD31^-CD326^-NG2^-PDPN^-FAP^-S100A4^+$. **(C)** Flow-activated cell sorting of the CAF population, gated on $CD45^-CD31^-EpCAM/CD326^-NG2^-PDGFR\alpha^+$.

**Supplementary Figure 3: CAF CM enhance the aggressive properties of triple-negative and epithelial cancer cells.**

**(A)** MTT assay analyzing the cell proliferation of 4T1**,** MDA-MB-231, and MCF-7 (n=4). $^{**}p < 0.01$ represents the significance level from a one-way ANOVA. **(B)** The addition of S100A4-enriched CAF CM compared to NF CM or cells only for wound percent closure of 4T1 and MDA-MB-231 cancer cells (n=4). $^{*}p < 0.05$, $^{**}p < 0.01$ represent the significance level from a one-way ANOVA. Scale bar: 400μm. **(C)** Images along with mean diameter and number of mammospheres of 4T1, MDA-MB-231, and MCF-7 (n=4). $^{**}p < 0.01$, $^{****}p < 0.0001$ represent the significance level from a one-way ANOVA. Scale bar: 120μm. Indicated (n) represents the number of independent experiments as biological replicates.

**Supplementary Figure 4: CAF cells enhance the aggressive properties when co-cultured with triple-negative cancer cells.**

**(A)** The addition of S100A4-enriched CAF cells compared to NF cells added or cells only for wound percent closure of GFP-tagged 4T1 and GFP-tagged MDA-MB-231 cancer cells (n=4). $^{***}p < 0.001$, $^{****}p < 0.0001$ represent the significance level from a one-way ANOVA. Scale bar: 400μm. **(B)** Images along with mean diameter and number of mammospheres of GFP-tagged 4T1 and GFP-tagged MDA-MB-231 cancer cells (n=4). $^{*}p < 0.05$, $^{****}p < 0.0001$ represent the significance level from a one-way ANOVA. Scale bar: 120μm. Indicated (n) represents the number of independent experiments as biological replicates.

**Supplementary Figure 5: CAF properties significantly attenuated by inhibition of S100A4 protein.**

**(A)** Immunofluorescence and cell counts of DAPI, S100A4, FAP, and PDPN-stained cells (n=5). *$p < 0.05$, ***$p < 0.001$ represent the significance level from a one-way ANOVA. Scale bar: 100µm. **(B)** Western blot analysis showing the expression of S100A4, FAP, Phospho-Ezrin, Ezrin, and PDPN in cell culture inhibition. The densitometric analyses compare the protein expression levels relative to β-actin (n = 4). *$p < 0.05$ represents the significance level from an unpaired *t*-test. Indicated (n) represents the number of independent experiments as biological replicates.

**Supplementary Figure 6: CAF CM inhibits NK cell cytotoxicity via co-culture.**

**(A)** Images of GFP-tagged Py230 spheroids co-cultured with spleen NK cells at 1:1 and 2:1 ratios in the presence and absence of S100A4-enriched CAF CM over 24 h (n=4). *$p < 0.05$, **$p < 0.01$, ****$p < 0.0001$ represents the significance level from a two-way ANOVA. Scale bar: 120µm. **(B)** Py230 and **(C)** NK cells metabolic assessment after conditioned media were transferred from $^{13}C_6$-glucose tracer-fed S100A4-enriched CAFs. **(D)** Heatmap from targeted metabolomics analysis of metabolites in NF and CAF samples (n=3). Indicated (n) represents the number of independent experiments as biological replicates.

**Supplementary Figure 7: CAF properties significantly attenuated by inhibition of BCAT1 and BCAT2 downstream targets.**

**(A)** ELISA MAX™ Deluxe Set Mouse IFN-γ was utilized to measure IFN-γ levels in cell culture supernatants of IL-12 and IL-15-activated spleen NK cells under various incubation conditions, utilizing BCAAs: valine, isoleucine, and leucine, and BCKAs: 3-Methyl-2-oxobutanoic acid (keto-valine), 3-Methyl-2-oxopentanoic acid sodium salt (keto-isoleucine), and α-Ketoisocaproic acid (keto-leucine) (*n* = 4). **$p < 0.01$, ***$p < 0.001$ represent the significance level from a one-way ANOVA. **(B)** Western blot analysis showing the expression of FOXM1 and SREBP1 in cell culture inhibition. The densitometric analyses compare the protein expression levels relative to β-actin (n = 4). *$p < 0.05$ represents the significance level from an unpaired *t*-test. Indicated (n) represents the number of independent experiments as biological replicates.
